# Aberrant laminin signaling drives melanocyte dedifferentiation and unveils a tractable therapeutic target in vitiligo

**DOI:** 10.1101/2025.04.11.648350

**Authors:** Fei Yang, Lingli Yang, Sylvia Lai, Masafumi Yokota, Yasutaka Kuroda, Takuo Yuki, Yoshito Takahashi, Tetsuya Sayo, Takeshi Namiki, Hiroyuki Goto, Sho Hiroyasu, Shigetoshi Sano, Shintaro Inoue, Yoichi Mizukami, Daisuke Tsuruta, Ichiro Katayama

## Abstract

Vitiligo is an acquired depigmenting skin disorder characterized by progressive melanocyte loss, but the cellular mechanisms driving this process remain unclear. Here, we identify melanocyte dedifferentiation as a central and reversible pathogenic mechanism in vitiligo. In healthy human skin, melanocytes reside within a basement membrane niche defined by dystroglycan-laminin-211 adhesion. In contrast, vitiligo lesions exhibit aberrant extracellular matrix remodeling, leading to an adhesion switch to integrin α3β1-laminin-332 interactions. This shift promotes melanocyte dedifferentiation *via* Rho-F-actin-dependent activation of Hippo and MAPK pathways, resulting in c-Jun-mediated transcriptional changes. Dedifferentiated melanocytes lose their pigment-producing identity and acquire neural crest-like features with multilineage potential. Importantly, this process is reversible. Pharmacological inhibition of involved pathways restores melanocyte differentiation and induces repigmentation in both vitiligo mouse models and *ex vivo* patient skin. Notably, JAK inhibitors also promote redifferentiation independently of immune modulation. These findings uncover melanocyte dedifferentiation as a fundamental driver of vitiligo and a tractable therapeutic target, offering new opportunities for therapeutic intervention.

## Introduction

Vitiligo is a common, acquired depigmenting disorder characterized by the progressive loss of functional melanocytes from the epidermis^1^. Although extensive studies have implicated autoimmune responses in disease initiation^2^, the precise mechanisms underlying melanocyte disappearance remain incompletely understood^3^. In addition to immune-mediated cytotoxicity, other factors, such as oxidative stress^4^, defective adhesion^5–7^, metabolic dysregulation^8^, and the accumulation of senescence-associated secretory phenotype factors^9^, have been proposed to contribute to melanocyte loss. However, whether melanocytes are entirely eliminated or instead persist in a nonfunctional or dedifferentiated state remains a fundamental and unresolved question with important clinical implications.

Melanocytes may persist in stable vitiligo lesions despite being morphologically altered, amelanotic, and undetectable by conventional melanocyte markers such as tyrosinase (TYR), tyrosinase-related protein 1 (TYRP1), dopachrome tautomerase (DCT), premelanosome protein (PMEL), melanoma antigen recognized by T-cells 1 (MART-1; also known as Melan-A), and melanocyte inducing transcription factor (MITF)^9–12^. These findings imply the existence of dedifferentiated or dormant melanocyte populations that evade standard detection, raising the possibility that changes in the local skin microenvironment, particularly at the basement membrane (BM), may play a critical role in regulating melanocyte identity and survival.

We previously reported structural abnormalities in the BM of vitiligo skin, including disorganized laminin deposition^13^. This finding is consistent with earlier histopathological studies that identified ultrastructural BM defects in vitiligo lesions^14–17^, collectively pointing to BM disruption as a potentially underappreciated factor in vitiligo pathogenesis^18–20^. Laminins, key components of the extracellular matrix (ECM), are known to regulate cell adhesion, polarity, and fate in various tissue types^21–23^. Given the well-established role of the ECM in defining tissue-specific niches and modulating cellular behavior, we hypothesized that BM alterations may contribute to melanocyte dysfunction in vitiligo by disrupting cell-matrix interactions.

Although the adhesion mechanisms of basal keratinocytes to the BM are well characterized, far less is known about how melanocytes anchor to and interact with their surrounding matrix. Although several adhesion molecules have been implicated in melanocyte-ECM binding^24–26^, the specific laminin isoforms involved, and their corresponding melanocyte receptors, remain poorly defined. Considering that laminins exhibit isoform-specific and spatially restricted expression patterns^22,27^, even subtle changes in BM composition may profoundly affect melanocyte adhesion, stability, and phenotype.

To address this gap, we systematically investigated the composition of laminin isoforms surrounding melanocytes in healthy and vitiligo skin and profiled their corresponding adhesion receptors. We found that in healthy skin, melanocytes adhere to laminin-211 *via* dystroglycan, forming a previously unrecognized adhesion architecture. In contrast, melanocytes in vitiligo lesions exhibited a striking adhesion switch to laminin-332 engagement *via* integrin α3β1. Functional analyses revealed that this switch promotes melanocyte dedifferentiation through Rho-F-actin-dependent activation of the Hippo and mitogen-activated protein kinase (MAPK) signaling pathways, ultimately leading to c-Jun-driven transcriptional changes and loss of pigment-producing identity.

Importantly, this dedifferentiated state was reversible. Pharmacological inhibition of key components in this pathway restored melanocyte differentiation and stimulated repigmentation in both vitiligo mouse models and patient-derived skin explants. These findings establish adhesion-dependent melanocyte dedifferentiation as a key pathogenic mechanism in vitiligo and highlight the BM microenvironment as a promising therapeutic target for restoring melanocyte function and skin pigmentation.

## Results

### Differential distribution of laminin isoforms in normal and vitiligo skin

First, we systematically examined the expression and distribution of laminin isoforms surrounding melanocytes in healthy and perilesional vitiligo skin. Immunofluorescence analysis revealed that in healthy skin, melanocytes were surrounded by laminin α2, β1, and γ1 (components of laminin-211), whereas laminin α3, β3, and γ2 (components of laminin-332) were selectively absent (Extended Data Fig. 1, Fig. 1a). In contrast, melanocytes in perilesional vitiligo skin exhibited prominent linear deposition of laminin α3, β3, and γ2, indicating a switch in the surrounding ECM composition (Fig. 1a). This observation was further supported by immuno-electron microscopy, which confirmed the absence of laminin α3 around melanocytes in healthy skin and its presence in vitiligo skin (Fig. 1b).

**Figure 1.**
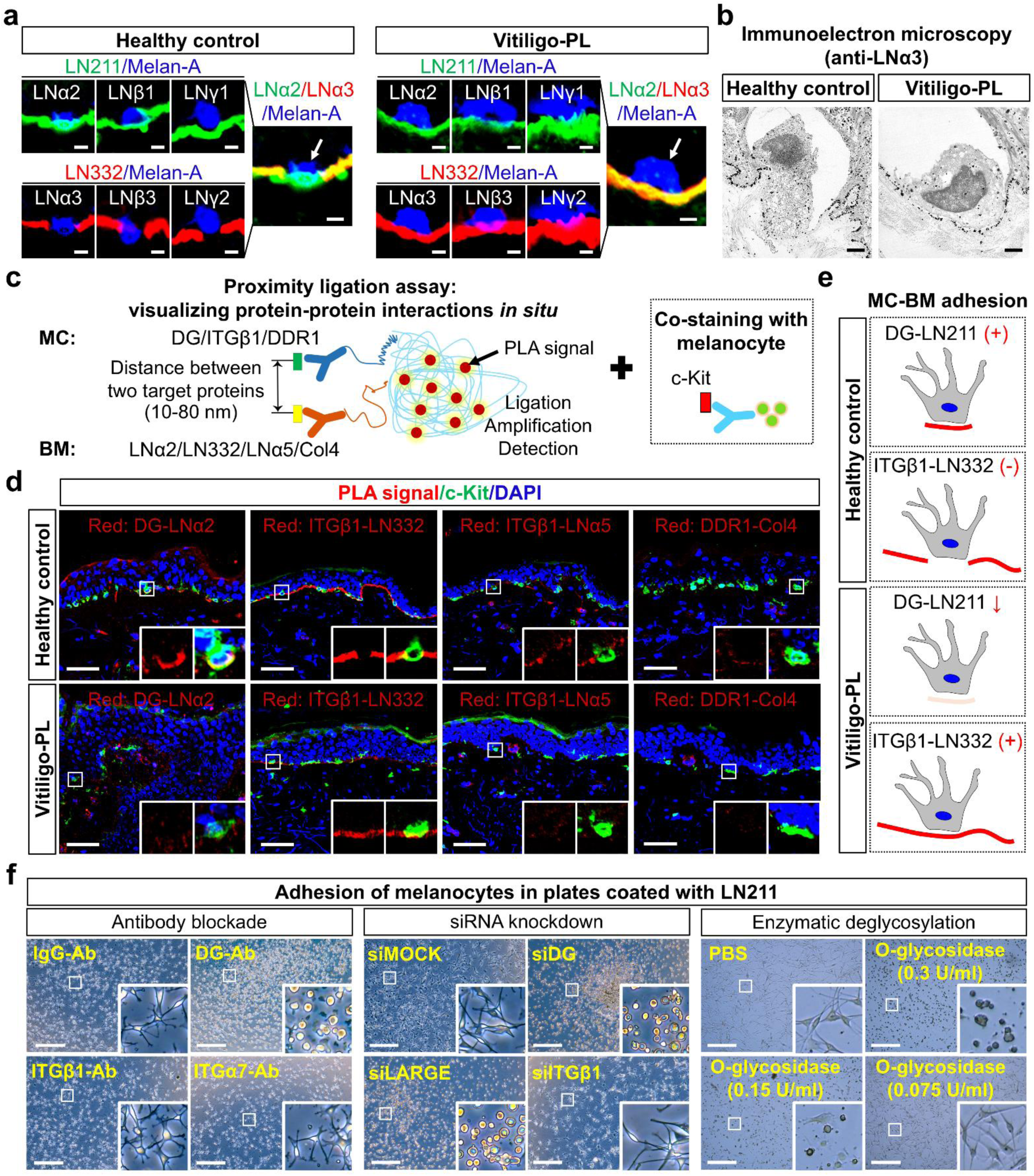
Distinct adhesion profiles of melanocytes in healthy and vitiliginous skin. **a**. Immunofluorescence analysis of laminin isoform distribution in healthy control skin and perilesional vitiligo skin (Vitiligo-PL). Laminin-211 (LN211) components include α2, β1, and γ1 chains; laminin-332 (LN332) components include α3, β3, and γ2 chains. Melanocytes are indicated by white arrows. Scale bar: 5 μm. **b**. Immuno-electron microscopy using anti-laminin α3 (LNα3) antibody reveals ultrastructural localization of laminin-332 around melanocytes in healthy and vitiligo-PL skin. Scale bar: 1 μm. **c**. Schematic of proximity ligation assay (PLA) for detecting protein-protein interactions between melanocyte surface receptors (e.g., DG, ITGβ1, DDR1) and basement membrane components (e.g., LNα2, LN332, LNα5, Col4). Melanocytes are costained using anti-c-Kit antibody. BM, basement membrane; MC, melanocyte. **d**. Representative PLA images showing red fluorescent signals (PLA puncta) indicating protein-protein interactions in healthy and vitiligo-PL skin. Costaining with c-Kit identifies melanocytes (green); nuclei are counterstained with 4′,6-diamidino-2-phenylindole (DAPI; blue). Scale bar: 50 μm. **e**. Proposed model of melanocyte-basement membrane (MC-BM) adhesion patterns in healthy and vitiligo-PL skin. **f**. Functional validation of DG-mediated melanocyte adhesion to LN211-coated plates using antibody blockade, small interfering RNA (siRNA) knockdown, and enzymatic deglycosylation. Scale bar: 500 μm. Ab, antibody; PBS, phosphate-buffered saline.

As integrins are currently known to serve as the main receptors for laminins^28^, we next examined the expression of integrin isoforms in normal epidermal melanocytes and detected α2, α5, αv, β1, β3, and low levels of β4 (Extended Data Fig. 2). To detect laminin-integrin interactions *in situ*, we performed the proximity ligation assay (PLA), an antibody-based technology that enables the visualization of protein-protein interactions (Fig. 1c), on skin tissue sections from healthy controls (Extended Data Fig. 3, Fig. 1d). In keratinocytes, the PLA signal validated previously reported interactions^29^, including integrin α6β4-laminin-332 (detected *via* integrin β4-laminin γ2), integrin α3β1-laminin-332 (detected *via* integrin β1-laminin γ2), and integrin α6β1-laminin-511 (detected *via* integrin β1-laminin α5) (Extended Data Fig. 3). However, in melanocytes, although integrin β1, β3, and β4 subunits were expressed (Extended Data Fig. 2), no detectable interaction with laminin-211 (identified *via* laminin α2), the major isoform surrounding melanocytes in healthy skin, was observed (Extended Data Fig. 3). Nevertheless, a weak interaction between integrin β1 and laminin γ1 was detectable in melanocytes (Extended Data Fig. 3); this binding was substantially weaker than the integrin-laminin interactions observed between keratinocytes and the BM. These findings suggest that in normal skin, melanocyte adhesion to the BM is unlikely to be primarily mediated by integrin-laminin-211 interactions.

Given prior reports that laminin-211 binds α-dystroglycan and integrin α7β1, particularly in muscle and neural tissues^27^, we investigated whether dystroglycan contributes to melanocyte adhesion. Using PLA, we observed strong dystroglycan-laminin-211 binding in normal melanocytes, whereas this interaction was markedly diminished in vitiligo melanocytes (Fig. 1d). This finding indicates that in healthy skin, melanocyte adhesion to laminin-211 is primarily mediated by dystroglycan.

In the basal epidermis, normal melanocytes were positioned in distinct laminin-332- and laminin α5- deficient niches along the BM (Extended Data Fig. 1), while in vitiligo skin, melanocytes were closely associated with BM regions enriched in laminin-332 (Extended Data Fig. 1, Fig. 1a). Together with prior evidence implicating discoidin domain receptor family, member 1 (DDR1)-collagen IV interactions in melanocyte adhesion^25^, we further examined these potential alternative adhesion pathways by PLA in healthy and perilesional vitiligo skin. In normal skin, melanocytes predominantly exhibited dystroglycan-laminin-211 binding, along with weak DDR1-collagen IV interactions. Notably, no laminin-332 or laminin α5 signals were detected at melanocyte-BM contact sites, resulting in melanocyte-specific gaps. In contrast, vitiligo melanocytes displayed robust integrin β1-laminin-332 binding, while other adhesion signals, including dystroglycan-laminin-211 and DDR1-collagen IV, were largely absent (Fig. 1d). These observations indicate that melanocyte adhesion mechanisms differ between healthy and vitiligo skin: normal melanocytes primarily rely on dystroglycan-laminin-211 interactions, whereas in vitiligo skin, adhesion is mediated predominantly through integrin β1-laminin-332 interactions (Fig. 1e).

To validate that dystroglycan-laminin-211 interaction is the predominant mechanism mediating melanocyte adhesion in healthy skin, we used antibody blockade, knockdown by small interfering RNA (siRNA), and enzymatic deglycosylation approaches. Blocking dystroglycan with specific antibodies significantly impaired melanocyte adhesion to laminin-211-coated surfaces, whereas no significant effect was seen with inhibition of integrin α7β1, the only other laminin-211 receptor identified to date aside from dystroglycan^27^. Similarly, siRNA-mediated knockdown of dystroglycan or its glycosylation regulator LARGE abolished adhesion to laminin-211. Enzymatic removal of glycosylation sites on dystroglycan also reduced melanocyte-laminin-211 binding in a dose-dependent manner (Fig. 1f). These findings uncover a clear molecular distinction between melanocyte adhesion mechanisms in healthy and vitiligo skin, reflecting a pathological reorganization of the BM microenvironment.

### Laminin-332-mediated adhesion shift induces melanocyte dedifferentiation

To investigate whether the altered adhesion environment observed in vitiligo skin affects melanocyte behavior, we cultured primary human epidermal melanocytes on laminin-332 or laminin-211 and compared their phenotypic and molecular characteristics. Interestingly, melanocytes on laminin-332 exhibited marked morphological changes, becoming flattened with reduced dendricity, in contrast to the highly branched and dendritic morphology observed on laminin-211 (Fig. 2a). Immunofluorescence analysis further revealed a profound decrease in expression of melanogenic proteins, including PMEL17 and TYRP1, under laminin-332 conditions (Fig. 2a).

**Figure 2.**
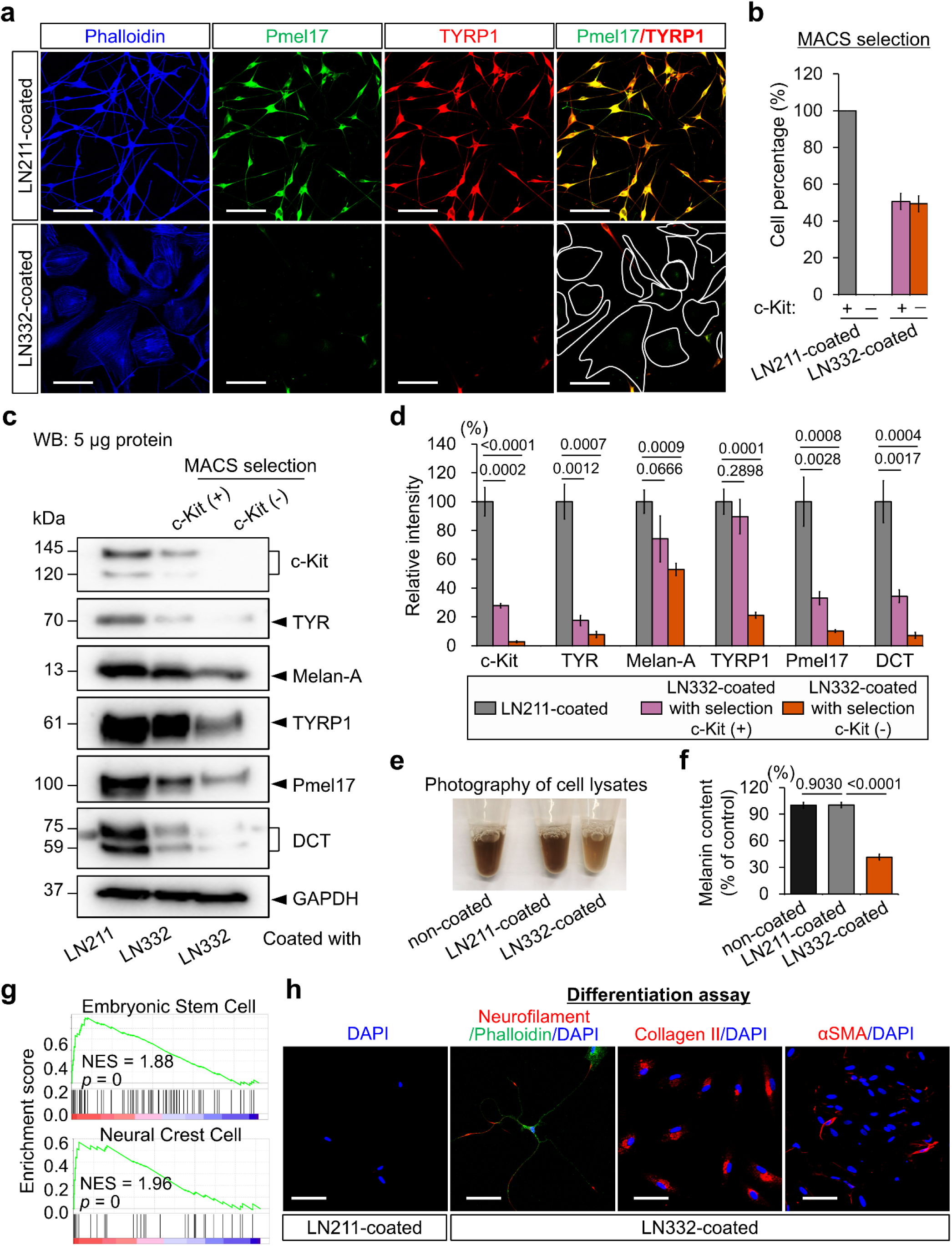
Laminin-332 induces dedifferentiation in human melanocytes. **a.** Morphological and phenotypic changes of melanocytes cultured on plates coated with laminin-211 (LN211) or laminin-332 (LN332). Cells were stained with phalloidin (F-actin, blue), PMEL17 (green), and TYRP1 (red). Merged images demonstrate reduced expression of melanocytic markers and altered cell morphology under LN332-coated conditions. Scale bar: 100 μm. **b.** Quantification of c-Kit-positive and c-Kit-negative melanocytes after magnetic-activated cell sorting (MACS) under LN211- and LN332-coated conditions. Data are shown as mean ± SD. Western blot (WB) analysis of melanocyte lineage markers in cells cultured on LN211 or LN332 substrates, with c-Kit-based MACS. Analyzed proteins include c-Kit, tyrosinase (TYR), Melan-A, TYRP1, PMEL17, and DCT. Glyceraldehyde 3-phosphate dehydrogenase (GAPDH) was used as a loading control. **d.** Quantification of protein expression levels shown in panel **c**. Data represent mean ± SD. Statistical significance was assessed by unpaired two-tailed Student’s t-test, and *P* values are indicated. **e.** Representative images of cell lysates from melanocyte pellets following culture under the indicated conditions. **f.** Quantification of melanin content from panel **e**, revealing a significant decrease in melanin levels in LN332-cultured cells. Data are shown as mean ± SD; *P* values (shown on graph) were calculated by one-way analysis of variance with Tukey’s *post hoc* test. **g.** Gene set enrichment analysis of melanocytes cultured on LN332, showing enrichment for embryonic stem cell and neural crest cell gene signatures, indicating loss of lineage identity. **h.** Assessment of multilineage differentiation potential in LN332-treated melanocytes. Cells were subjected to lineage-specific induction protocols and immunostained for neurofilament (neuronal), collagen II (chondrogenic), and alpha-smooth muscle actin (α-SMA; myogenic) markers. Scale bar: 100 μm. DAPI, 4′,6-diamidino-2-phenylindole.

To assess the effect of laminin-332 on melanocyte phenotype, we performed magnetic-activated cell sorting (MACS) based on expression of c-Kit, a cell surface receptor that serves as a well-established marker of the melanocytic lineage. Compared with melanocytes cultured on laminin-211, cells exposed to laminin-332 exhibited a markedly increased c-Kit-negative population (Fig. 2b). Although c-Kit is not a definitive marker of differentiation status, the loss of its expression may indicate a change in differentiation status, suggesting that laminin-332 induces cellular reprogramming. Western blot analysis revealed markedly reduced expression of key melanogenic proteins, including DCT, TYR, PMEL17, TYRP1, and Melan-A, in both c-Kit-negative and c-Kit-positive subsets from laninin-332-treated cells compared with those cultured on laminin-211 (Fig. 2c, d). Consistent with these molecular changes, melanin content was significantly decreased in laminin-332-treated cultures (Fig. 2e, f). This phenotype was consistently observed across five independent primary melanocyte lines (Extended Data Fig. 4), indicating that the response to laminin-332 is reproducible and not limited to a specific cell line.

The dramatic changes in melanocyte morphology, marker expression, and pigment production prompted us to investigate whether laminin-332 induces a dedifferentiated cell state. Transcriptomic profiling followed by gene set enrichment analysis (GSEA) revealed significant upregulation of gene signatures associated with embryonic stem cells and neural crest cells in laminin-332-treated melanocytes compared with those cultured on laminin-211 (Fig. 2g), indicating the acquisition of a more plastic, progenitor-like state. To test the functional significance of this reprogramming, we assessed the multilineage potential of laminin-332-treated melanocytes in lineage-specific differentiation assays. Upon exposure to appropriate differentiation media, these cells exhibited the capacity to transdifferentiate into neurofilament-positive neurons, collagen II-producing chondrocytes, and alpha-smooth muscle actin-positive muscle-like cells (Fig. 2h), confirming their dedifferentiated and multipotent phenotype. These results demonstrate that altered interactions with laminin-332 induce a dedifferentiated melanocyte state, characterized by molecular and phenotypic changes and reduced melanogenic capacity.

### Molecular and signaling mechanisms underlying laminin-332-induced melanocyte dedifferentiation and modulation by external stimuli

To investigate the molecular mechanisms underlying laminin-332-induced melanocyte dedifferentiation, we performed bulk RNA sequencing of melanocytes cultured on laminin-332 or laminin-211. Laminin-332-treated cells exhibited a distinct transcriptional profile, with a large number of differentially expressed genes compared with laminin-211-treated controls (Fig. 3a). Melanogenic genes such as *MLANA*, *PMEL*, and *TYR* were significantly downregulated, consistent with the observed reduction in pigment production (Fig. 2). Conversely, several genes associated with the Hippo signaling pathway, including *TGFBI*, *CCN1*, *DKK1*, *CCN2*, and *THBS1*, were markedly upregulated (Fig. 3a). Pathway enrichment analysis further revealed significant activation of multiple signaling cascades, including the Hippo, PI3K (phosphatidylinositol 4,5-bisphosphate 3-kinase catalytic subunit beta isoform)-Akt (protein kinase B), and MAPK pathways (Fig. 3b). Enrichment of cytoskeleton-related pathways may also account for the cell flattening and morphological changes observed in laminin-332-treated melanocytes (Fig. 2a).

**Figure 3.**
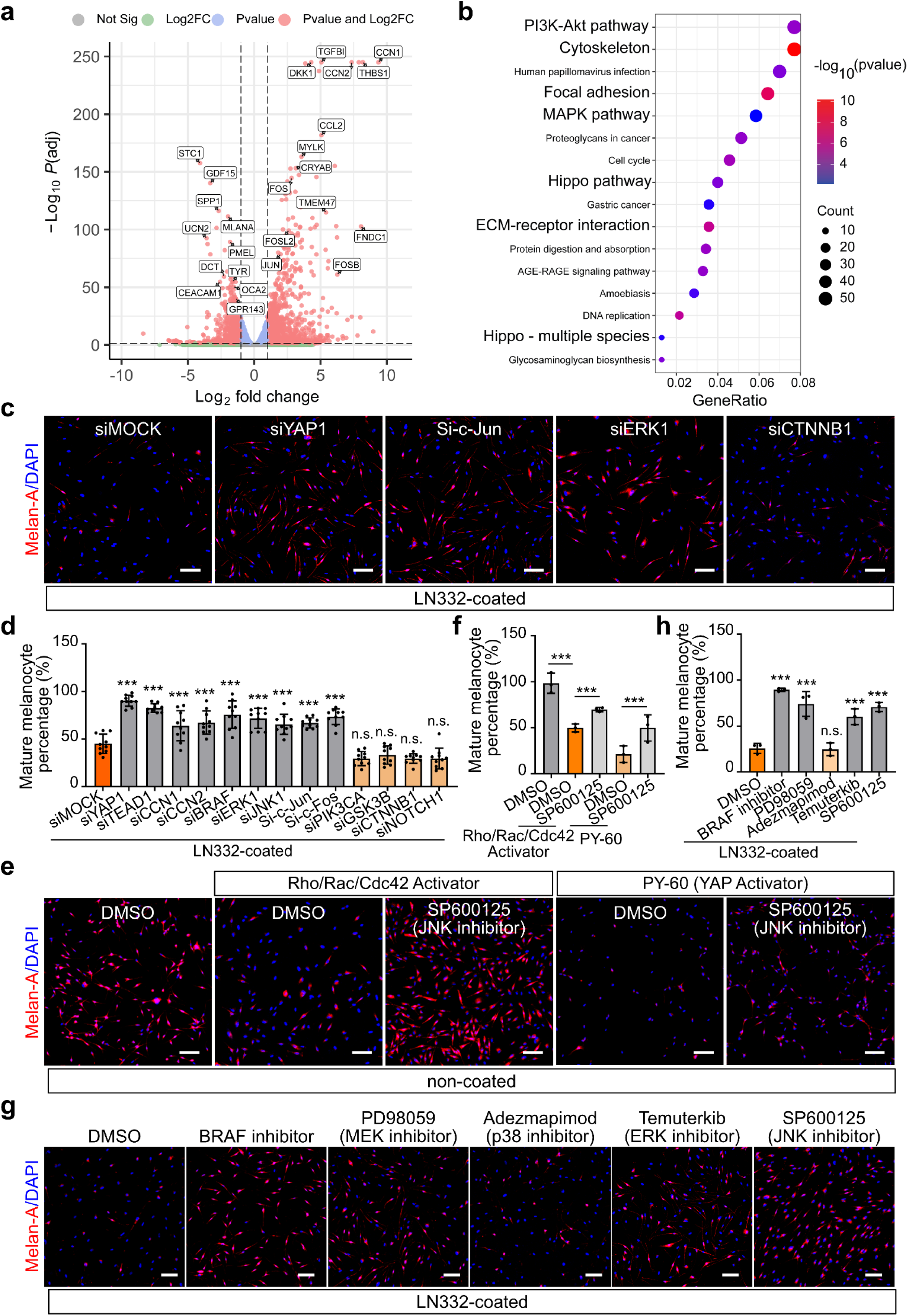
Molecular mechanisms underlying laminin-332-induced dedifferentiation in melanocytes. **a.** Volcano plot showing differentially expressed genes in melanocytes cultured on laminin-332 (LN332) compared with laminin-211 (LN211), based on bulk RNA-seq analysis. Thresholds: |log_2_ fold change| > 1 and false discovery rate < 0.05. **b.** Gene ontology enrichment analysis of upregulated pathways in LN332-treated melanocytes. Enriched signaling pathways include PI3K-Akt, MAPK, focal adhesion, and Hippo signaling. **c.** Representative immunofluorescence images of melanocytes after small interfering RNA (siRNA)-mediated knockdown of YAP1, c-Jun, ERK1, and CTNNB1 (β-catenin). Cells were stained for Melan-A (red) and nuclei (4′,6-diamidino-2-phenylindole, DAPI; blue). Scale bar: 100 μm. **d.** Quantification of mature melanocyte populations after gene knockdown. Data represent mean ± SD. Statistical significance was assessed using one-way analysis of variance with *post hoc* testing. \*\*\**P* < 0.001; n.s., not significant. **e.** Rescue of melanocyte differentiation on noncoated plates *via* cotreatment with pathway activators and inhibitors. Representative images showing the effect of JNK inhibition (SP600125) on cells treated with Rho/Rac/Cdc42 activator or PY-60 (YAP activator). Scale bar: 100 μm. DMSO, dimethyl sulfoxide. **f-h.** Quantification of melanocyte differentiation under various pathway-modulator treatments corresponding to panels **e** and **g**. Data are shown as mean ± SD; \*\*\**P* < 0.001; n.s., not significant. **g.** Rescue of melanocyte differentiation on LN332-coated plates using inhibitors targeting the MAPK pathway, including BRAF, MEK, p38, ERK, and JNK. Cells were stained for Melan-A (red) and nuclei (DAPI, blue). Scale bar: 100 μm.\

To functionally validate the pathways implicated by transcriptomic analysis, we systematically silenced key components of the Hippo, MAPK, and PI3K-Akt pathways in melanocytes cultured on laminin-332. Small interfering RNA-mediated knockdown of Hippo pathway effectors (transcriptional coactivator YAP1; transcriptional enhancer factor TEF-1 [also known as TEAD1]; CCN family member 1 [CCN1]; CCN family member 2 [CCN2]) and MAPK components (serine/threonine-protein kinase B-raf [BRAF]; mitogen-activated protein kinase 3 [MAPK3, also known as ERK1]; JNK1/MAPK8-associated membrane protein [JNK1]) significantly increased the proportion of mature melanocytes, indicating suppression of dedifferentiation (Fig. 3c, d). Similarly, knockdown of activator protein 1 (AP-1) transcription factors c-Jun and c-Fos restored melanocyte differentiation (Fig. 3c, d). In contrast, knockdown of phosphatidylinositol 4,5-bisphosphate 3-kinase catalytic subunit alpha isoform (PIK3CA), glycogen synthase kinase-3 beta (GSK3B), catenin beta-1 (CTNNB1), or neurogenic locus notch homolog protein 1 (NOTCH1) did not significantly affect the dedifferentiated phenotype, suggesting that the PI3K-Akt pathway and other canonical differentiation-related regulators do not play a major role in laminin-332-induced dedifferentiation.

Given that integrin signaling can activate both Hippo and MAPK cascades *via* Rho GTPase-mediated actin remodeling^30,31^, we tested whether direct cytoskeletal activation was sufficient to induce a dedifferentiation phenotype comparable to that observed with laminin-332 treatment. Strikingly, treatment with a Rho/Rac/Cdc42 activator, which promotes actin cytoskeleton reorganization, induced melanocyte dedifferentiation even in the absence of laminin-332, closely resembling the phenotype observed under laminin-332 treatment (Fig. 3e, f). These findings align with prior studies showing that both Hippo and MAPK pathways can activate AP-1 transcriptional programs^32,33^. Western blot analysis further confirmed that dedifferentiated melanocytes exhibited increased expression of cytoskeletal regulators (RhoA, Rac1/2/3, Cdc42, F-actin), Hippo effectors (YAP, WW domain-containing transcription regulator protein 1 [TAZ], CCN1, CCN2), MAPK signaling components (p-ERK, p-JNK), and AP-1 transcription factors, with c-Jun showing the strongest induction (5.6-fold increase) (Extended Data Fig. 5a-d; quantified in Extended Data Fig. 5e). These results demonstrate that c-Jun acts as a key downstream effector in laminin-332-induced melanocyte dedifferentiation.

To further validate the role of c-Jun, we pharmacologically inhibited its upstream activator JNK using SP600125, a selective JNK inhibitor. This intervention effectively blocked dedifferentiation induced by both Rho/Rac/Cdc42 and YAP activation (Fig. 3e, f). In addition, inhibitors targeting the MAPK cascade revealed that BRAF, MEK, and especially JNK inhibition suppressed dedifferentiation, whereas p38 inhibition had no significant effect (Fig. 3g, h). These data are consistent with transcriptional network analyses showing that JNK and ERK, but not p38, strongly regulate c-Jun expression and activity^34^, highlighting c-Jun as the key convergence point of the Rho-F-actin-Hippo-MAPK axis in melanocyte dedifferentiation.

We next investigated whether melanocyte dedifferentiation could be modulated by vitiligo-associated extrinsic factors. Treatment of melanocytes with inflammatory cytokines (tumor necrosis factor alpha [TNFα], interferon beta [IFNβ], interferon gamma [IFNγ], interleukin 1 beta [IL1β], interleukin 6 [IL6]) or an oxidative stress inducer (hydrogen peroxide [H_2_O_2_]), all of which have been implicated in vitiligo pathogenesis^35,36^, induced dedifferentiation, as evidenced by loss of melanocyte markers (Extended Data Fig. 6a). However, the dedifferentiation response to these stimuli was less pronounced than that induced by laminin-332 (Extended Data Fig. 6a), suggesting that diverse stress-related signals can trigger melanocyte plasticity with varying efficiencies.

Interestingly, several JAK inhibitors, including ruxolitinib, delgocitinib, baricitinib, abrocitinib, and tofacitinib, markedly suppressed laminin-332-induced melanocyte dedifferentiation (Extended Data Fig. 6b). Given that these agents are typically used to suppress immune activation, our findings suggest an additional, previously unrecognized mechanism by which JAK inhibition may directly promote melanocyte differentiation, independent of immune modulation.

In light of our previous finding that the epidermal growth factor receptor (EGFR) inhibitor gefitinib improved repigmentation in vitiligo animal models^37^, we further tested whether other tyrosine kinase inhibitors could affect melanocyte dedifferentiation. Inhibitors targeting EGFR, vascular endothelial growth factor receptor 2, insulin-like growth factor 1 receptor (IGF1R), angiopoietin-1 receptor (Tie2), c-RET, and ALK tyrosine kinase receptor (ALK) all attenuated laminin-332-induced dedifferentiation, with particularly strong effects observed for EGFR, IGF-1R, c-RET, and ALK inhibitors (Extended Data Fig. 6c).

Numerous studies have previously linked tyrosine kinase signaling, reactive oxygen species, and proinflammatory cytokines to the activation of MAPK and AP-1 pathways^33,38–40^. Collectively, our findings demonstrate that melanocyte dedifferentiation is governed by an integrin-Rho-F-actin-Hippo/MAPK-c-Jun signaling axis, and that this process is further modulated by extrinsic stimuli, including tyrosine kinase signaling, oxidative stress, and cytokines (Extended Data Fig. 6d).

### Rho-F-actin-Hippo/MAPK-c-Jun signaling pathway regulates melanocyte differentiation across *in vitro*, *in vivo*, and *ex vivo* systems

To confirm the functional relevance of the signaling pathways identified in Fig. 3, we examined the effects of modulating Rho-F-actin, Hippo, and MAPK (JNK) signaling across three model systems: cultured primary melanocytes (Fig. 4a), K14-Kitl transgenic mice (Fig. 4b), and *ex vivo* human skin explants (Fig. 4c).

**Figure 4.**
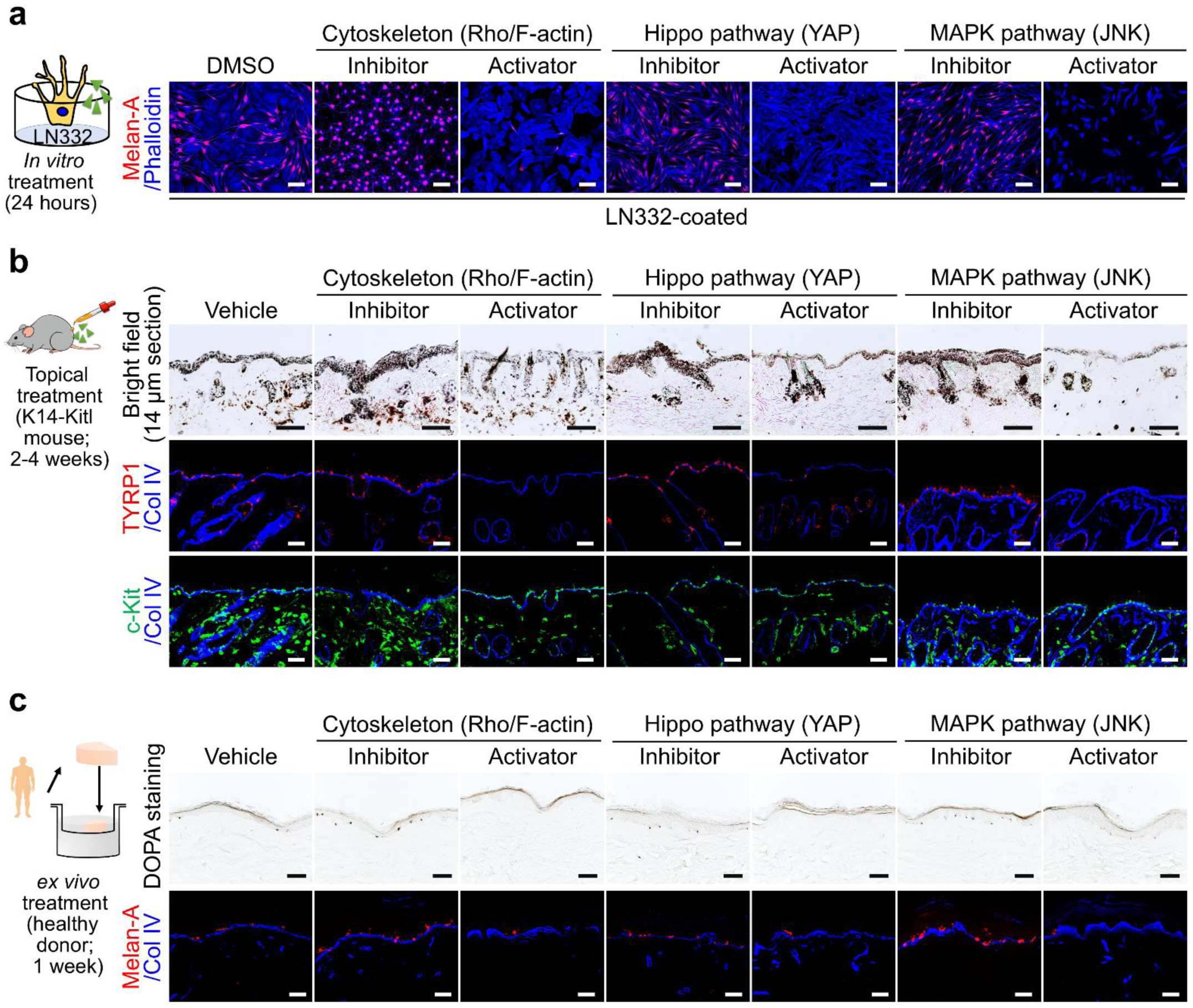
Validation of Rho/Hippo/MAPK pathway involvement in melanocyte differentiation across multiple models. **a.** *In vitro* effects of pathway modulation on melanocyte differentiation. Human melanocytes were cultured on plates coated with laminin-332 (LN332) and treated with specific inhibitors or activators targeting the Rho/F-actin cytoskeleton, Hippo (YAP), or MAPK (JNK) pathways for 24 hours. Cells were stained with anti-Melan-A (red) and phalloidin (F-actin; blue). Scale bar: 100 μm. DMSO, dimethyl sulfoxide. **b.** *In vivo* validation using K14-Kitl mice subjected to topical treatment with pathway modulators for 2-4 weeks. Top panels: bright-field images of dorsal skin sections showing pigmentation status. Middle panels: immunofluorescence staining for mature melanocytes using anti-TYRP1 (red) and anti-collagen IV (blue). Bottom panels: staining for immature melanocytes using anti-c-Kit (green) and anti-collagen IV (blue). c-Kit single-positive cells located in the epidermis were defined as immature melanocytes. Scale bar: 50 μm (immunofluorescence), 100 μm (bright-field). **c.** *Ex vivo* human skin explant model treated with pathway modulators for 1 week. Top panels: 3,4-dihydroxyphenylalanine (DOPA) staining visualizing functionally mature melanocytes. Bottom panels: immunofluorescence staining for mature melanocytes (Melan-A, red; collagen IV, blue). Scale bar: 100 μm (DOPA), 50 μm (immunofluorescence).

In melanocyte cultures maintained on laminin-332, approximately 50% of cells retained a mature phenotype, as indicated by Melan-A expression. Treatment with Rho/F-actin, YAP, or JNK inhibitors markedly increased the proportion of Melan-A-positive melanocytes and markedly reduced the population of dedifferentiated phalloidin-positive cells, effectively restoring a fully differentiated phenotype (Fig. 4a). In contrast, treatment with activators of these pathways led to a pronounced reduction in mature melanocytes and a corresponding increase in dedifferentiated cells. These findings support the conclusion that inhibition of Rho-Hippo/MAPK signaling promotes melanocyte differentiation, while activation of these pathways drives dedifferentiation, consistent with the mechanistic findings described above (Fig. 3).

We next assessed these effects *in vivo* using K14-Kitl transgenic mice, which exhibit melanocyte retention in the dorsal epidermis. Two-week topical applications of Rho, YAP, or JNK inhibitors enhanced epidermal pigmentation, as observed by bright-field microscopy (Fig. 4b). Immunofluorescence analysis revealed increased numbers of mature melanocytes (TYRP1⁺/c-Kit⁺) in the basal epidermis. Conversely, 4-week treatment with corresponding pathway activators led to hypopigmentation and decreased numbers of mature melanocytes (TYRP1⁺/c-Kit⁺) in the basal epidermis, further confirming that these signaling pathways regulate melanocyte differentiation status *in vivo*.

To evaluate the relevance of these findings in human tissue, we applied the same modulators to healthy adult human skin explants. After 1 week of treatment, Rho, YAP, or JNK inhibition increased the number of functional melanocytes, as evidenced by increased 3,4-dihydroxyphenylalanine (DOPA) staining and Melan-A expression in the basal layer (Fig. 4c). In contrast, pathway activators reduced both DOPA⁺ and Melan-A⁺ melanocyte populations.

Together, these results demonstrate that Rho-F-actin-Hippo/MAPK-c-Jun pathways act as central regulators of melanocyte differentiation across multiple biological systems and ultimately determine pigmentation outcomes in skin.

### Melanocyte dedifferentiation in patients with vitiligo

Given our earlier observation that melanocytes in vitiligo skin are in direct contact with laminin-332 (Fig. 1), we next investigated whether the dedifferentiation mechanisms identified *in vitro* are also present in patients’ tissue. We first assessed melanogenic activity in perilesional skin using the DOPA reaction. Compared with healthy controls, melanocytes in perilesional vitiligo skin showed markedly reduced DOPA staining (Fig. 5a), demonstrating a reduction in pigment-producing capacity.

**Figure 5.**
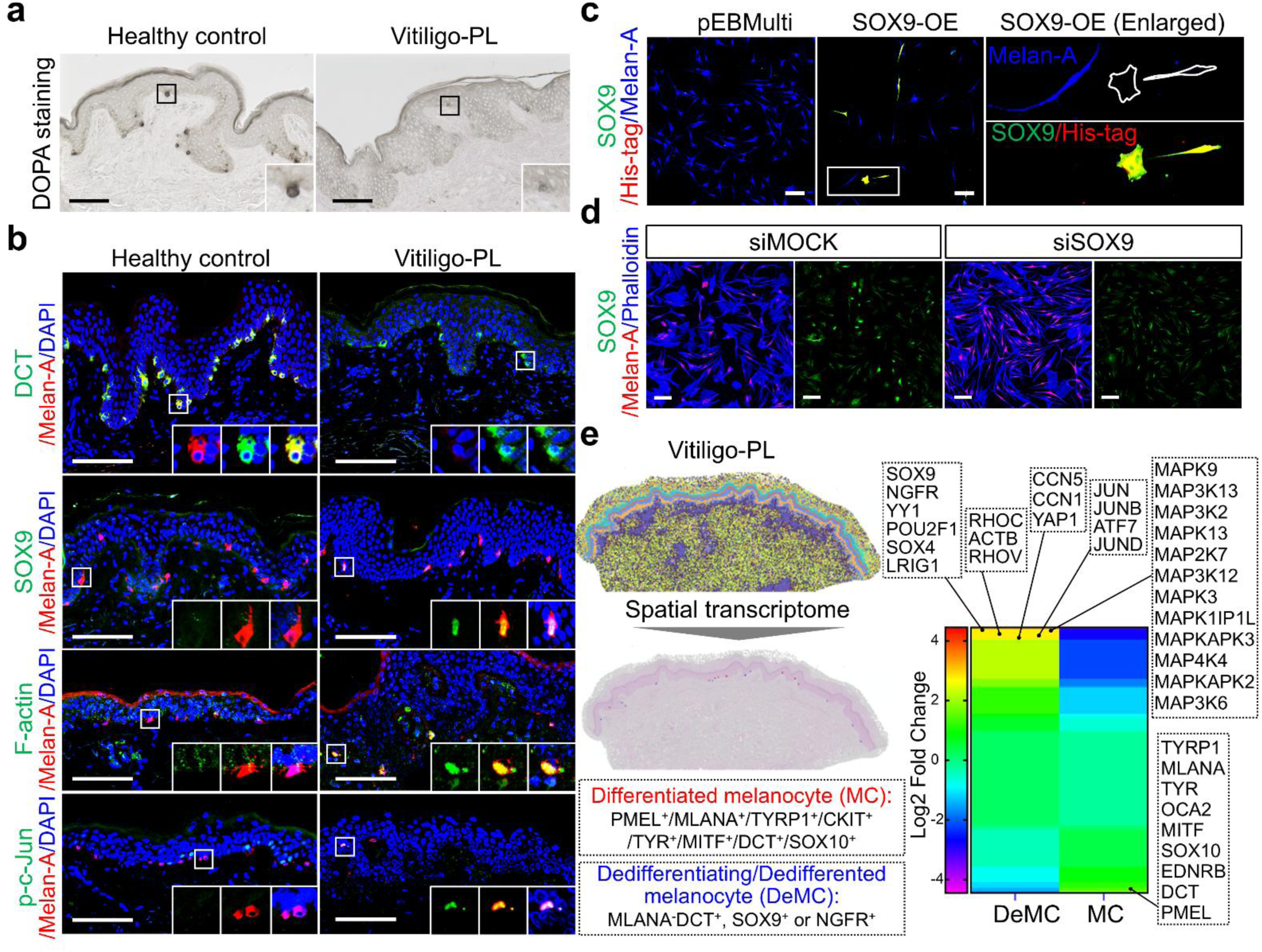
Evidence of melanocyte dedifferentiation in skin from patients with vitiligo. **a.** 3,4-dihydroxyphenylalanine (DOPA) staining of healthy control and perilesional vitiligo (Vitiligo-PL) skin reveals reduced melanocyte function in vitiliginous areas. Scale bar: 100 μm. **b.** Immunofluorescence analysis of melanocyte markers in healthy and Vitiligo-PL skin. Staining includes DCT (green), SOX9 (green), F-actin (green), p-c-Jun (green), Melan-A (red), and 4′,6-diamidino-2-phenylindole (DAPI; blue). Insets highlight melanocytes with altered marker expression patterns in Vitiligo-PL samples. Scale bar: 50 μm. **c.** Overexpression (OE) of SOX9 in cultured melanocytes (SOX9-OE) enhances melanocyte dedifferentiation. Cells were stained for SOX9 (green), His-tag (red), and Melan-A (blue). Enlarged view highlights SOX9/His-tag positive cells with low Melan-A expression. Scale bar: 100 μm. **d.** SOX9 knockdown by small interfering RNA (siRNA; siSOX9) promotes melanocyte differentiation in cultured melanocytes. Cells were stained for SOX9 (green), Melan-A (red), and phalloidin (blue). Scale bar: 100 μm. **e.** Spatial transcriptomics of perilesional vitiligo skin identifies two melanocyte populations: mature differentiated melanocytes (MC; PMEL⁺/MLANA⁺/TYRP1⁺/MITF⁺/DCT⁺/SOX10⁺) and dedifferentiating or dedifferentiated melanocytes (DeMC; MLANA⁻/DCT^+^, SOX9⁺ or NGFR⁺). Gene expression heatmap (log_2_ fold change) highlights upregulation of dedifferentiation-related genes (*SOX9*, *NGFR*, *YAP1*, *RHOV*, *JUN* family) in DeMC, and loss of pigment genes (*TYRP1*, *MLANA*, *MITF*) compared with MC.

To identify reliable markers for dedifferentiated melanocytes in patient samples, we first compared laminin-332-treated melanocytes with cultured melanoblasts. Both cell types lacked Melan-A expression, but although melanoblasts typically retained early melanocytic markers such as c-Kit and DCT^41^, laminin-332-treated cells showed heterogeneous or absent expression of these markers, suggesting a dedifferentiation state beyond the melanoblast stage. Notably, both melanoblast and laminin-332-induced dedifferentiated cells exhibited strong expression of neural crest-associated markers SOX9 and tumor necrosis factor receptor superfamily member 16 (NGFR), consistent with their developmental origin^42^ (Extended Data Fig. 7). Further immunophenotyping of laminin-332-treated melanocytes revealed a spectrum of differentiation states: from fully differentiated cells (Melan-A⁺/DCT⁺ or c-Kit⁺, SOX9⁻/NGFR⁻) (Extended Data Fig. 8, cell type #1), to intermediate dedifferentiating states (e.g., SOX9⁺ or NGFR⁺ with partial retention of Melan-A) (Extended Data Fig. 8, cell type #2), to fully dedifferentiated cells lacking all melanocytic markers but expressing SOX9 and/or NGFR (e.g., NGFR⁺/SOX9⁺) (Extended Data Fig. 8, cell type #3). These observations provided a rationale for using SOX9 and NGFR as reliable markers for dedifferentiated melanocytes in tissue analysis.

Although c-Kit and NGFR were undetectable in most perilesional melanocytes (Extended Data Fig. 9), the combined assessment of DCT/SOX9 and Melan-A reliably identified dedifferentiating cells (Fig. 5b). Specifically, dedifferentiated melanocytes were characterized either by a DCT-positive/Melan-A-negative phenotype or by concurrent SOX9/Melan-A expression (Fig. 5b). These dedifferentiated cells often expressed elevated F-actin and phosphorylated c-Jun (Fig. 5b), consistent with the cytoskeletal and transcriptional changes observed under laminin-332–induced conditions *in vitro* (Fig. 3; Extended Data Fig. 5).

Previous studies have shown that SOX9 is expressed in melanoblasts and downregulated during melanocyte differentiation^43^. SOX9 also antagonizes SOX10^44^, a master regulator of melanocyte differentiation *via* MITF activation^45^, and plays a critical role in precursor fate determination in the hair follicle niche^46^. Increased SOX9 expression has been reported in vitiligo melanocytes^47^. To test the functional relevance of SOX9 in melanocyte dedifferentiation, we overexpressed SOX9 in cultured melanocytes using an episomal pEBMulti vector. SOX9 overexpression, confirmed by His-tag colabeling, led to a marked decrease in Melan-A expression (Fig. 5c), indicating loss of a mature melanocyte phenotype.

We next examined whether SOX9 is required for laminin-332–induced dedifferentiation. In melanocytes cultured on laminin-332, siRNA-mediated knockdown of SOX9 (siSOX9) markedly suppressed the dedifferentiation phenotype such that the proportion of Melan-A⁺ mature melanocytes significantly increased compared with control (siMOCK)-transfected cells (Fig. 5d). These results demonstrate that SOX9 is necessary for the dedifferentiation program triggered by laminin-332. As c-Jun has previously been shown to transcriptionally activate SOX9^48^, our findings suggest that SOX9 functions downstream of the Rho-F-actin-Hippo/MAPK-c-Jun axis to drive phenotypic reprogramming of melanocytes in vitiligo.

Spatial transcriptome analysis of perilesional vitiliginous skin provided further molecular evidence for melanocyte dedifferentiation. To classify melanocyte states, we first identified melanocyte-like cells based on the presence of translucent cytoplasm on hematoxylin and eosin (H&E)-stained sections. Cells within this morphological category were further classified as differentiated melanocytes (MCs) if they expressed canonical melanogenic markers such as Melan-A or PMEL. In contrast, dedifferentiated or dedifferentiating melanocytes (DeMCs) were defined as translucent H&E-positive cells that lacked Melan-A/PMEL expression but showed positivity for DCT, SOX9, or NGFR.

Gene expression profiling revealed that DeMCs exhibited upregulation of genes associated with stemness (e.g., *SOX9*, *NGFR*, *YY1*, *POU2F1*, *SOX4*, *LRIG1*), cytoskeletal remodeling (e.g., *RHOC*, *ACTB*, *RHOV*), Hippo signaling components (e.g., *CCN5*, *CCN1*, *YAP1*), AP-1 family transcription factors (e.g., *JUN*, *JUNB*, *ATF7*, *JUND*), and MAPK pathway kinases (e.g., *MAPK9*, *MAP3K13*, *MAP3K2*, *MAPK13*) (Fig. 5e). In contrast, MCs predominantly expressed melanogenic genes such as *TYRP1*, *MLANA*, *TYR*, and *MITF* (Fig. 5e). The transcriptional signatures of DeMCs in patient skin closely mirrored those observed in laminin-332-treated melanocytes *in vitro*, supporting the presence of a conserved dedifferentiation program.

### Pharmacological reversal of melanocyte dedifferentiation restores pigmentation across multiple vitiligo models

To explore the therapeutic potential of targeting melanocyte dedifferentiation, we tested the efficacy of pharmacological inhibitors across multiple vitiligo model systems.

We first developed a three-dimensional (3D) human skin model with defined laminin-332 exposure, allowing precise modulation of the melanocyte microenvironment. Laminin-332 coating induced a concentration-dependent loss of pigmentation (Fig. 6a), generating sharply demarcated depigmented zones (Fig. 6b) that recapitulate the lesion morphology observed in vitiligo. Immunostaining confirmed that laminin-332 treatment decreased the number of Melan-A-positive mature melanocytes (Fig. 6c) while increasing DCT-single-positive dedifferentiating melanocytes (Extended Data Fig. 10), consistent with a dedifferentiation phenotype. Treatment with the JNK inhibitor SP600125 effectively reversed these changes, restoring pigmentation and increasing the proportion of mature melanocytes (Fig. 6c).

**Figure 6.**
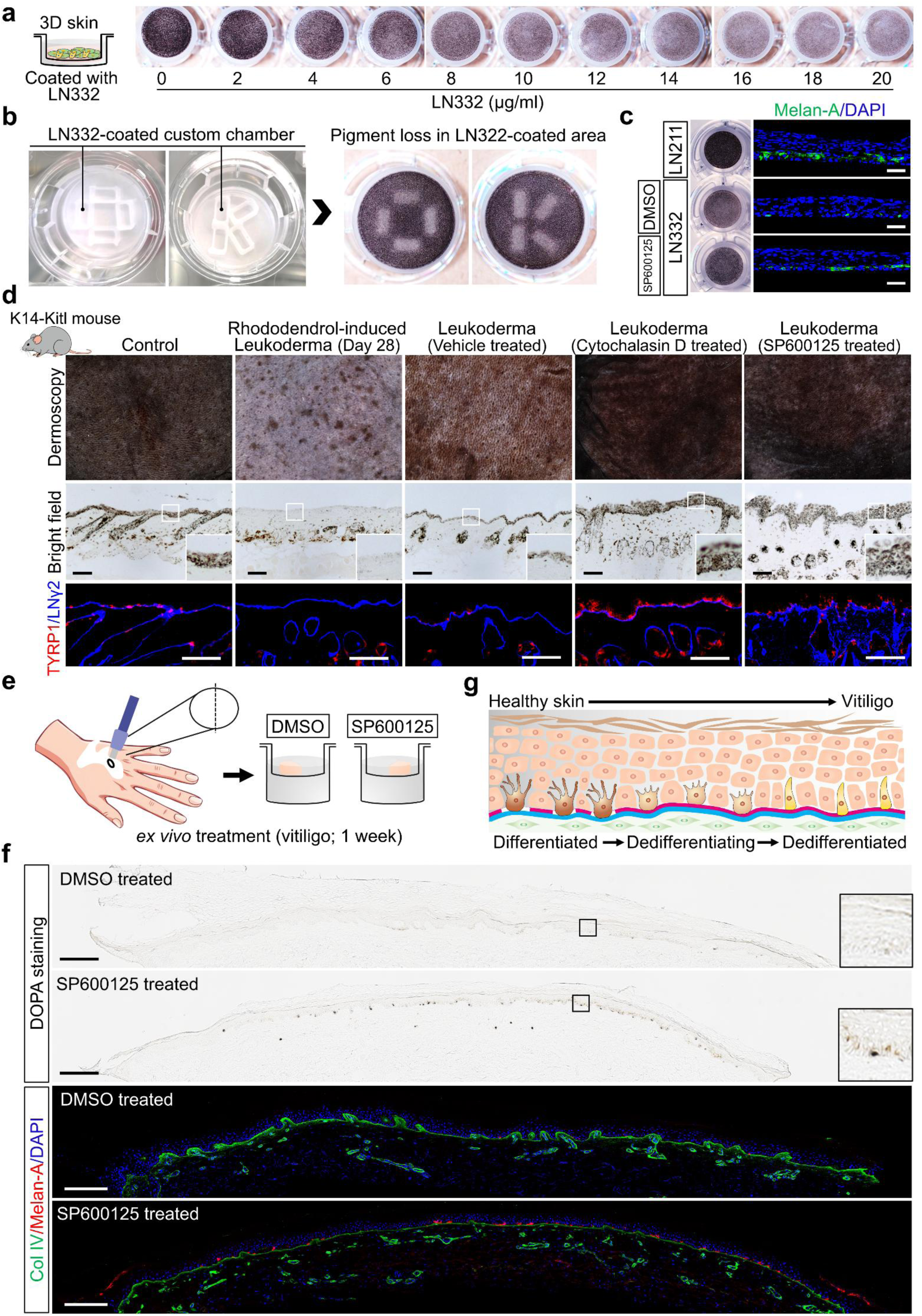
Therapeutic efficacy of dedifferentiation-targeted therapy in vitiligo models. **a.** Dose-dependent effects of laminin-332 (LN332) on pigmentation in a three-dimensional (3D) reconstructed human skin model. LN332 was applied at increasing concentrations (0-20 μg/mL), resulting in progressive pigment loss in a concentration-dependent manner. **b.** Localized pigment loss induced by LN332 in a custom-designed culture chamber. Melanocytes cultured on LN332-coated areas exhibited patterned depigmentation. **c.** Rescue of pigmentation in the 3D skin model following treatment with the dedifferentiation inhibitor SP600125 (a JNK inhibitor). Cells were stained for Melan-A (green) and nuclei (4′,6-diamidino-2-phenylindole, DAPI; blue). Scale bar: 50 μm. DMSO, dimethyl sulfoxide. **d.** *In vivo* therapeutic effect of dedifferentiation inhibitors in a rhododendrol-induced leukoderma mouse model (K14-Kitl mice). Skin pigmentation was assessed by dermoscopy and bright-field imaging. Immunofluorescence staining for TYRP1 (red) and LNγ2 (blue) demonstrated restored melanocyte differentiation and pigmentation following SP600125 treatment. Scale bar: 100 μm. **e.** *Ex vivo* treatment with SP600125 for 1 week of perilesional vitiligo skin from patients. **f.** 3,4-dihydroxyphenylalanine (DOPA) staining and immunofluorescence analysis of treated human skin explants. SP600125 restored functional melanocytes, as indicated by increased pigmentation and Melan-A expression (red). Collagen IV is shown in green, and DAPI in blue. Scale bar: 200 μm. **g.** Schematic illustration of melanocyte states in healthy versus vitiligo skin, illustrating the transition from differentiated to dedifferentiated states and the therapeutic reversal by dedifferentiation inhibition.

To validate these findings *in vivo*, we used a rhododendrol-induced depigmentation model in K14-Kitl mice^49^. Rhododendrol application resulted in progressive pigment loss in both fur and skin (Extended Data Fig. 11a, b), with prominent melanocyte dedifferentiation at intermediate concentrations (10%) (Extended Data Fig. 11c). In this rhododendrol-induced depigmentation model, both cytochalasin D (an actin polymerization inhibitor) and SP600125 significantly restored pigmentation and increased TYRP1-positive melanocytes in the epidermis (Fig. 6d), further supporting a central role for cytoskeletal and JNK signaling in melanocyte dedifferentiation.

We next evaluated the therapeutic potential of dedifferentiation inhibition using *ex vivo* lesional skin explants from patients with vitiligo. After 1 week of SP600125 treatment, DOPA staining revealed a marked increase in DOPA-positive melanocytes compared with dimethyl sulfoxide (DMSO)-treated controls, indicating restoration of melanogenic activity (Fig. 6f, top panels). Immunofluorescence analysis further showed a significant increase in Melan-A-positive melanocytes in SP600125-treated samples, with these cells localized along the BM as defined by collagen IV staining (Fig. 6f, bottom panels). In untreated explants, many melanocytes were DCT-positive but lacked Melan-A expression, consistent with a dedifferentiated phenotype. In contrast, SP600125 treatment induced re-expression of Melan-A in these DCT-positive cells (Extended Data Fig. 12), supporting the reversal of the dedifferentiated state and restoration of a mature melanocyte identity.

Together, these findings demonstrate that the inhibition of melanocyte dedifferentiation restores pigmentation across 3D skin equivalents, mouse models, and patient-derived skin explants, supporting its central role in vitiligo pathogenesis and suggesting that dedifferentiation reversal may represent a potential therapeutic strategy for vitiligo (Fig. 6g).

## Discussion

Following a “clinical-to-basic-to-clinical” research approach, this study identifies and elucidates the molecular mechanism of melanocyte dedifferentiation in vitiligo and demonstrates its reversibility across human tissue and disease models. In the present study, we uncovered a previously unrecognized mechanism contributing to melanocyte loss in vitiligo. Rather than a complete disappearance of melanocytes, cells in lesional skin exhibit a dedifferentiated phenotype. In healthy human skin, melanocytes are located within specialized BM niches characterized by the presence of laminin-211 and the absence of laminin-332. In contrast, vitiligo skin displays pathological BM remodeling, with aberrant accumulation of laminin-332 around melanocytes. This shift in laminin composition disrupts cell-matrix interactions and triggers intracellular signaling cascades. Through systematic pathway analysis, we demonstrate that laminin-332 binding to integrin β1 initiates Rho-dependent cytoskeletal remodeling, leading to activation of the Hippo and MAPK pathways, ultimately resulting in c-Jun activation. Functionally, this signaling suppresses melanocyte differentiation markers, reduces melanin synthesis, and induces a transcriptional program associated with neural crest-like progenitor cells. Notably, this dedifferentiation process is reversible. Pharmacological inhibition of key signaling restored pigmentation and melanogenic marker expression in 3D skin equivalents, a rhododendrol-induced leukoderma mouse model, and *ex vivo* patient-derived skin explants.

To understand how this laminin-332-deficient niche is maintained in healthy skin and disrupted in vitiligo, we investigated the role of melanocyte-derived matrix-degrading enzymes. In normal human skin, melanocytes expressed high levels of the membrane-type matrix metalloproteinases MT1-MMP (MMP14) and MT4-MMP (MMP17), as confirmed by immunostaining and validated in multiple skin-derived cell lines (Extended Data Fig. 13a-d). Functional assays further demonstrated that melanocytes actively degrade laminin-332, forming clear pericellular clearance zones; this activity was significantly impaired upon knockdown of either MT1-MMP or MT4-MMP (Extended Data Fig. 13e). In vitiligo, epidermal melanocytes exhibited reduced MT4-MMP expression, accompanied by abnormal accumulation of laminin-332 (Extended Data Fig. 14a-b). These findings are consistent with our previous observations of aberrant laminin-332 distribution and BM disorganization in vitiligo skin^13^. Although the mechanisms underlying MT4-MMP downregulation remain to be clarified, decreased expression or functional abnormalities of MT4-MMP in melanocytes diminished their ability to degrade laminin-332, thereby destabilizing this specialized niche. Moreover, other laminin-332-degrading enzymes^50^, including MMP2, MMP9, MMP3, neutrophil elastase, cathepsin G, plasmin, and others, may release laminin-332 fragments that accumulate around melanocytes and further disrupt niche integrity. In vitiligo, we previously observed an enrichment of MMP2 near the BM^13^. These concurrent alterations may accelerate the damage to the laminin-332-deficient niche, ultimately promoting melanocyte dedifferentiation in vitiligo. Additionally, mouse melanocytes lacked expression of both MT1-MMP and MT4-MMP (Extended Data Fig. 14c), limiting the translational utility of mouse models for studying melanocyte-matrix interactions and ECM remodeling in human skin.

In addition to ECM remodeling, alterations in melanocyte adhesion receptors contribute to niche instability. In healthy skin, melanocytes primarily adhere to the BM *via* dystroglycan-laminin-211 interactions (Fig. 1d). In vitiligo, this adhesion is markedly reduced (Fig. 1d), likely due to decreased dystroglycan expression in perilesional melanocytes (Extended Data Fig. 15a). Given that dystroglycan is a known substrate of MMP2^51^, and our previous studies have shown elevated MMP2 levels in patients with vitiligo^13^, this reduction may be attributed to enhanced proteolytic degradation by MMP2. We also found that exposure to laminin-332 downregulates dystroglycan expression in melanocytes (Extended Data Fig. 15b), suggesting a potential feedback loop. Reduced dystroglycan weakens laminin-211 binding, facilitating compensatory engagement with laminin-332, which further promotes dedifferentiation and suppresses dystroglycan expression. These findings establish dystroglycan as a critical adhesion receptor for melanocytes. Interestingly, dystroglycan defects also lead to melanocyte detachment in congenital muscular dystrophy (CMD)^52^, although patients with CMD do not display vitiligo-like depigmentation, possibly due to the absence of dedifferentiation-inducing cues such as laminin-332 accumulation. Given dystroglycan’s conserved roles in muscle and nervous systems^53^, and the shared neural crest origin of melanocytes^54^, our findings suggest that melanocytes may serve as a useful model for investigating CMD-associated neurodevelopmental abnormalities.

The dedifferentiation phenotype described here has important implications for vitiligo pathogenesis. Flattened melanocytes observed in vitiliginous skin and culture have been previously noted^55–57^, and some have been shown to regain function upon oxidative stress relief^10^. Our findings provide molecular evidence that melanocytes in lesional vitiligo skin are not completely absent, but rather persist in a dedifferentiated, nonpigmented state. Although we identified candidate markers, including c-Kit, DCT, NGFR, and SOX9, the heterogeneity in their expression, along with the presence of fully dedifferentiated, marker-negative cells, suggests that additional work is needed to define robust and universal markers for detecting dedifferentiated melanocytes in human tissue. This concept of reversible melanocyte dedifferentiation is further supported by recent work in hair follicle biology. A 2023 study by Sun et al.^50^ demonstrated that melanocyte stem cells within the hair follicle niche dynamically transition between stem- and transit-amplifying states through reversible dedifferentiation in response to local cues such as WNT signaling. These findings align with our observations in vitiligo, suggesting that melanocyte fate plasticity, whether in hair follicles or the interfollicular epidermis, may be broadly governed by niche-dependent and reversible dedifferentiation programs.

Mechanistically, we showed that melanocyte dedifferentiation was mediated by the integrin α3β1-Rho-F- actin-Hippo/MAPK-c-Jun signaling upon laminin-332 binding. Notably, JNK inhibition with SP600125 reversed this process, restoring melanocyte morphology and pigmentation. Although inhibitors targeting Rho, YAP, and JNK were effective in our models, their current clinical applications are limited to oncology, indicating the need for further development of melanocyte-targeted compounds suitable for vitiligo treatment. Interestingly, JAK inhibitors enhanced melanocyte differentiation independent of their immunomodulatory effects, possibly *via* STAT3-mediated regulation of AP-1 components c-Jun/c-Fos^58^. This mechanism further supports the clinical application of JAK inhibitors, although more targeted agents may offer promising efficacy and safety in the future.

Cellular differentiation and dedifferentiation are fundamental processes in development, regeneration, and disease. Although dedifferentiation has been extensively studied in cultured cells, particularly in induced pluripotent stem cells^59^, it occurs rarely in adult human tissues and typically under severe stress or injury^60,61^. Our findings establish melanocytes as a model for stress-induced dedifferentiation in skin. In studies of ocular lineage differentiation from induced pluripotent stem cells, laminin-211 has been shown to direct differentiation toward the neural crest lineage, whereas laminin-332 directs differentiation toward the epithelial lineage^62^. This finding aligns with our observation that laminin-211 supports melanocyte differentiation, whereas laminin-332 promotes dedifferentiation (Fig. 2). Furthermore, mice engineered to express laminin-211 in the BM show a human-like melanocyte localization in the basal layer^63^, indicating laminin-211’s essential role in niche structure and melanocyte retention.

Taken together, our study demonstrates that the pathological microenvironment in vitiligo could trigger melanocyte dedifferentiation through Rho-F-actin-dependent activation of Hippo and MAPK pathways, ultimately leading to c-Jun activation (Fig. 3 and Extended Data Fig. 16). Consistent with previous reports, Hippo signaling is known to regulate epithelial tissue reprogramming^64^ and cooperate with paired box protein Pax-3to modulate MITF expression in neural crest cells, thereby influencing melanocyte fate specification^65^. In addition to the core pathway identified, our data suggests that other microenvironmental factors, including reactive oxygen species, proinflammatory cytokines, and other tyrosine kinase inhibitors, also contribute to melanocyte differentiation (Extended Data Fig. 6). Although individual cytokines and tyrosine kinase inhibitors exert modest effects, their combined influence *in vivo* may be substantially amplified. These findings indicate the importance of targeting both intrinsic signaling and extrinsic niche components in therapeutic strategies. The identification of melanocyte dedifferentiation as a reversible process not only offers new therapeutic opportunities for vitiligo (Fig. 6) but also raises the possibility of modulating skin pigmentation under physiological conditions (Fig. 4), with potential applications in cosmetic dermatology and pigmentation disorders. This work represents a significant advance in understanding vitiligo pathogenesis and highlights promising new therapeutic strategies. Future research should focus on identifying more specific markers for dedifferentiated melanocytes and on developing safe, selective targeted therapies.

## Methods

### Human skin specimens

Frozen biopsies were obtained from lesional and perilesional skin of well-defined patients with nonsegmental vitiligo (n = 15, ten patients in a progressive state and five patients in a stable state) (Supplementary Table 1) and from corresponding sites on healthy donors (n = 7). The progressive state of vitiligo was defined by the development of new lesions or the extension of pre-existing lesions in the previous 6 months, and the stable state was defined by no increase in the size of existing lesions and an absence of new lesions in the previous year. Written informed consent was obtained from all participants prior to study inclusion. The study was approved by the Medical Ethics Committee of Osaka Metropolitan University (No. 4152). All procedures involving human subjects were in accordance with the Helsinki Declaration of 1975, as revised in 1983.

### Cell culture

Primary human melanocytes, including HEMn-DP (#C2025C, Thermo Fisher Scientific, Waltham, MA, USA), HEMn-MP (#C1025C, Thermo Fisher Scientific), and other human epidermal melanocytes (#LMC, #LC1C1022, and #M4C0760) obtained from Kurabo Co. (Osaka, Japan), were cultured in medium 254 (Cascade Biologics, Invitrogen, Waltham, MA,, USA). Melanoblasts (36011-12, Celprogen, Torrance, CA, USA) were maintained in Human Melanoblast Stem Cell Serum Free Media (#M36011-12, Celprogen) using precoated flasks (#E36011-12-T75, Celprogen). All cell lines were incubated at 37 °C in a humidified atmosphere containing 5% CO_2_. For laminin-coating experiments, culture plates were treated overnight at 4 °C with either 20 μg/mL laminin-211 (#LN211-02, BioLamina, Sundbyberg, Sweden) or 20 μg/mL laminin-332 (#LN332-02, BioLamina).

### Antibody neutralization experiments

Melanocytes were preincubated with one of the following antibodies at a final concentration of 10 μg/mL for 30 minutes: rabbit IgG control (#ab172730, Abcam, Cambridge, UK), mouse IgG control (#ab37355, Abcam), anti-dystroglycan (#ab234587, Abcam), anti-integrin β1 (#PA5-29606, Thermo Fisher Scientific), and anti-integrin α7 (#ab195959, Abcam). After pretreatment, cells were seeded onto laminin-211–coated coverglass (#C018001, Matsunami Glass Ind., Ltd., Osaka, Japan) and incubated for 4–6 hours before evaluating cell adhesion.

### Melanin content measurement

Harvested cells were solubilized in Solvable solution (6NE9100, Perkin Elmer, Waltham, MA, USA) at 80 °C for 30 minutes. The melanin content of each sample was then quantified by measuring its absorbance at 405 nm using a spectrophotometer. A standard curve generated with a synthetic melanin standard (M8631, Sigma-Aldrich, St. Louis, MO, USA) was used to calculate the melanin concentration in each sample.

### Multilineage differentiation

To assess differentiation potential, laminin-332–treated, dedifferentiated melanocytes were cultured in lineage-specific induction media. For neural differentiation, cells were maintained in Mesenchymal Stem Cell Neurogenic Differentiation Medium (#C-28015, PromoCell, Heidelberg, Germany). For chondrogenic differentiation, the MesenCult-ACF Chondrogenic Differentiation Medium (#ST-05455, Veritas, Tokyo, Japan) was used. For mesodermal differentiation, cells were cultured in a 1:1 mixture of Dulbecco’s Modified Eagle’s Medium (DMEM; #D5921, Sigma-Aldrich) and F-12 (#N4888, Sigma-Aldrich), supplemented with 1× nonessential amino acid solution (#139-15651, FUJIFILM Wako Pure Chemical Corporation, Osaka, Japan), 60 pM recombinant human TGF-beta 1 protein (#240-B-010/CF, R&D Systems, Minneapolis, MN, USA), 10% fetal calf serum (#10270106, Thermo Fisher Scientific), and 1× penicillin-streptomycin solution (#168-23191, FUJIFILM Wako Pure Chemical Corporation). Differentiation was evaluated after 1–4 weeks using appropriate lineage-specific markers.

### MACS cell sorting

c-Kit-positive and c-Kit-negative cell populations were isolated using MACS. Sorting was performed using a MiniMACS Starting Kit (#130-090-312, Miltenyi Biotec, Bergisch Gladbach, Germany) and a CD117 MicroBead Kit (#130-091-332, Miltenyi Biotec), following the manufacturer’s instructions.

### 3D skin model

Three-dimensional skin models containing a dermal compartment were constructed in accordance with our previously established protocol^13^. Briefly, human dermal fibroblasts were embedded in an acidic collagen solution (KKN-IAC-50, Koken Co. Ltd, Tokyo, Japan). Following a 48-hour gel contraction period, melanocytes or melanoblasts were coseeded with keratinocytes onto the contracted dermal equivalent. For spatially controlled laminin-332 treatment, specific regions were defined using culture inserts (#80209, ibidi GmbH, Gräfelfing, Germany). After laminin-332 coating and removal of the culture inserts, melanocytes and keratinocytes were coseeded to form the epidermal layer. These 3D skin models were subsequently maintained at the air-liquid interface for 10 days.

### Mouse experiments

Eight-week-old male K14-Kitl mice (#RBRC00694, RIKEN BioResource Research Center, Tsukuba, Japan) were used for all *in vivo* experiments. Rhododendrol (4-(4-hydroxyphenyl)-2-butanol) was generously provided by Kanebo Cosmetics Inc. (Tokyo, Japan). A stock solution of 20% (w/v) rhododendrol was prepared in a 50:50 (v/v) ethanol/sesame oil mixture as the vehicle. To establish a leukoderma mouse model, mice were treated topically with 100 μL of 20% (w/v) rhododendrol on the dorsal skin once daily for 1 month.

This induction period was followed by a 2-week drug intervention phase. To specifically examine melanocyte dedifferentiation, additional mice received a single topical application of rhododendrol 24 hours before tissue collection and analysis. Specifically, 100 μL of a 5%, 10%, or 15% (w/v) rhododendrol solution was applied to the dorsal skin of each mouse. All animal experiments were conducted in accordance with protocols approved by the Institutional Animal Care and Use Committee (Approval No. 23018).

### Immunofluorescence and immunohistochemistry

Frozen skin sections (6-μm thick) were fixed in 4% paraformaldehyde (163-20145, FUJIFILM Wako Pure Chemical Corporation) for 15 minutes at room temperature, followed by blocking with 5% bovine serum albumin for 30 minutes. Sections were then incubated with primary antibodies (Supplementary Table 2), diluted 1:100 in blocking solution, overnight at 4 °C. Following primary antibody incubation, sections were washed three times for 5 minutes each with phosphate-buffered saline (PBS) containing 0.05% Tween 20 (161-24801, FUJIFILM Wako Pure Chemical Corporation). Appropriate secondary antibodies (Supplementary Table 3), diluted 1:1000 in blocking solution, were then applied for 1 hour at room temperature in the dark. Sections were subsequently washed three times with 0.05% Tween 20 in PBS for 5 minutes per wash. Nuclei were counterstained with 4′,6-diamidino-2-phenylindole (DAPI; 340-07971, FUJIFILM Wako Pure Chemical Corporation). Slides were imaged using an LSM 710 confocal laser scanning microscope (ZEISS Group, Oberkochen, Germany), and images were acquired with ZEN 2012 software (ZEISS Group). Quantification of positively stained cells and mean fluorescence intensity was performed using ImageJ software (National Institutes of Health, https://imagej.net/ij/index.html).

### Immuno-electron microscopy

Skin tissues were fixed in 4% paraformaldehyde (163-20145, FUJIFILM Wako Pure Chemical Corporation) for 30 minutes at room temperature, and then washed three times in PBS (5 minutes each).

Tissues were then cryoprotected by immersion in 30% sucrose at 4 °C overnight and subsequently embedded in OCT compound (#45833, Funakoshi Co., Ltd., Tokyo, Japan). Frozen sections (7 μm thick) were prepared and permeabilized in 0.25% saponin (#47036, MilliporeSigma, Burlington, MA, USA) for 30 minutes. Sections were blocked for 30 minutes in PBS containing 0.1% saponin, 10% bovine serum albumin (#014- 15151, FUJIFILM Wako Pure Chemical Corporation), 10% normal goat serum (#16210072, Thermo Fisher Scientific), and 0.1% gelatin from cold water fish skin (#G7041, MilliporeSigma). Samples were then incubated with the primary antibody against laminin α3 (#FDV-0024, Funakoshi) diluted in blocking buffer overnight at 4 °C. Following several washes, the sections were incubated with nanogold-conjugated anti-mouse IgG (1.4 nm in diameter; 1:200; #2001, Nanoprobes, Yaphank, NY, USA) in blocking solution for 2 hours at room temperature. The nanogold signal was enhanced using the GoldEnhance EM kit (#2113, Nanoprobes) for 4 minutes at room temperature. Samples were postfixed in 1% osmium tetroxide (#157- 01141, FUJIFILM Wako Pure Chemical Corporation), dehydrated through a graded ethanol series, and embedded in TAAB EPON 812 embedding resin (#342-2, Nisshin EM, Tokyo, Japan). Ultrathin sections (70 nm) were cut, stained with uranyl acetate and lead citrate, and imaged using a Talos F200C G2 transmission electron microscope (Thermo Fisher Scientific) equipped with a CCD-based camera system (Advanced Microscopy Techniques, Woburn, MA, USA).

### Proximity ligation assay

Frozen human skin sections were blocked with Duolink blocking solution (#DUO82007, MilliporeSigma) for 30 minutes at 37 °C. The sections were then incubated overnight at 4 °C with primary antibodies (diluted 1:100). After this incubation, the sections were washed in Duolink *In Situ* Wash Buffer A (#DUO82046, MilliporeSigma) and subsequently incubated with Duolink *In Situ* PLA Probes (Anti-Mouse PLUS, #DUO92001; Anti-Rabbit MINUS, #DUO92005; MilliporeSigma) for 90 minutes at 37 °C. This step was followed by ligation and signal amplification using the Duolink *In Situ* Detection Reagents Red kit (#DUO92008, MilliporeSigma). For costaining of melanocytes, an anti-c-Kit antibody (#AF332, R&D Systems) was included along with the other primary antibodies during the overnight incubation. During the PLA probe incubation step, a corresponding Alexa Fluor Plus 488–conjugated donkey anti-goat IgG (#A32814, Thermo Fisher Scientific) was added to detect the c-Kit primary antibody. This approach allowed the simultaneous visualization of melanocytes in the tissue sections during the PLA probe incubation step. After a final wash in Duolink *In Situ* Wash Buffer B (#DUO82048, MilliporeSigma) for 30 minutes, the sections were mounted with Duolink *In Situ* Mounting Medium containing DAPI (#DUO82040, MilliporeSigma) to counterstain nuclei. The slides were then examined under an LSM 710 confocal laser scanning microscope (ZEISS Group), and fluorescent images were captured using ZEN 2012 software (ZEISS Group).

### RNA sequencing and analysis

RNA sequencing and library preparation were performed by Takara Bio Inc. (Shiga, Japan) using the SMART-Seq v4 Ultra Low Input RNA protocol. Sequencing was conducted on a NovaSeq 6000 platform with NovaSeq Control Software v1.7.5, Real Time Analysis v3.4.4, bcl2fastq2 v2.20, and DRAGEN Bio-IT Platform v3.9.3 (Illumina, San Diego, CA, USA). Each experimental group included three biological replicates, with a mean sequencing depth of approximately 55.08 million reads per sample. Reads were aligned to the human genome (GRCh38.primary_assembly.genome.fa.gz) using GENCODE annotation (v39; gencode.v39.primary_assembly.annotation.gtf.gz). Differential gene expression analysis was conducted using DESeq2, and gene ontology enrichment analysis was performed with SRplot^66^. Gene set enrichment analysis (GSEA) was carried out using GSEA v4.3.3 in accordance with the official GSEA User Guide, with gene sets derived from the c2.cp.wikipathways.v2024.1.Hs.symbols collection.

### Spatial transcriptomics

Spatial transcriptomic analysis was performed using the Visium HD Spatial Gene Expression platform (10x Genomics, Pleasanton, CA, USA) at the Yamaguchi University Institute of Gene Research. A formalin-fixed, paraffin-embedded (FFPE) skin tissue block from a perilesional area of patients with nonsegmental vitiligo was analyzed. From each FFPE block, two consecutive 5-μm sections were cut and directly mounted onto Visium HD capture slides. RNA was extracted from adjacent sections to assess integrity and purity; only samples meeting internal quality thresholds for degradation proceeded to library preparation. After placement, the tissue sections underwent H&E staining. A 6.5-mm × 6.5-mm capture area containing 2-μm high-density spots was used to spatially profile approximately 18,000 genes at near single-cell resolution. Downstream data processing, including quality control, gene expression mapping, and clustering analysis, was performed using Loupe Browser (10x Genomics), enabling spatially resolved transcriptomic profiling of the vitiligo skin tissue.

### Western blot analysis

Protein extraction from cell pellets and the subsequent Western blot analysis were performed as described in our previous study^67^. In each assay, 5 μg of extracted protein was used. Primary antibodies were applied at the dilutions specified in Supplementary Table 4, with the anti-glyceraldehyde 3-phosphate dehydrogenase antibody serving as the internal loading control. Signal intensities of the immunoreactive bands were quantified using ImageJ densitometry software (National Institutes of Health, https://imagej.net/ij/index.html).

### DOPA reaction

Frozen skin sections were fixed in 1% glutaraldehyde for 1 hour and then washed 2 to 3 times with PBS. The sections were then incubated at 37 °C for 2 hours in a 0.1% L-DOPA solution (#043-30563, FUJIFILM Wako Pure Chemical Corporation) to facilitate melanin synthesis, as L-DOPA serves as a substrate for tyrosinase activity. After incubation, the sections were washed 2 to 3 times with PBS and postfixed with glutaraldehyde (#071-02031, FUJIFILM Wako Pure Chemical Corporation). Subsequently, the sections were washed, dehydrated through an ethanol series, and mounted for microscopic observation.

### siRNA knockdown

Human epidermal melanocytes were transfected with siRNAs using Lipofectamine RNAiMAX transfection reagent (#13778030, Thermo Fisher Scientific) in accordance with the manufacturer’s instructions. Each siRNA was used at a final concentration of 30 nM. Six hours after transfection, the medium was replaced with fresh growth medium. Knockdown efficiency for each target gene was confirmed by quantitative real-time polymerase chain reaction (qPCR) on extracted RNA. A nontargeting siRNA (siMOCK; #AM4611, Thermo Fisher Scientific) was used as a negative control. Target-specific siRNAs (Thermo Fisher Scientific Silencer Select siRNAs) were used to knock down the genes listed in Supplementary Table 5. All siRNA reagents were purchased from Thermo Fisher Scientific.

### Glycosidase digestion and cell adhesion assay

Melanocytes were seeded on laminin-211–coated coverglass (#C018001, Matsunami Glass Ind., Ltd.). Glycosidase digestion was performed in Opti-MEM medium (#31985070, Thermo Fisher Scientific) at 37 °C. To remove O-linked oligosaccharides from α-dystroglycan on the cell surface, cells were first incubated with neuraminidase (0.1-0.5 U/mL; #P0720S, New England Biolabs, Ipswich, MA, USA) for 2 hours. This neuraminidase treatment cleaved terminal sialic acid residues, thereby exposing the underlying O-linked oligosaccharide chains on α-dystroglycan. After the 2-hour incubation, the neuraminidase-containing solution was aspirated, and cells were gently washed with PBS. Subsequently, cells were incubated with O-glycosidase (0.05-0.5 U/mL; #P0733S, New England Biolabs) in fresh Opti-MEM medium for another 2 hours to remove the exposed oligosaccharides. Controls were treated with Opti-MEM-containing enzyme buffer alone under identical conditions. Cell adhesion to laminin-211-coated coverslips was evaluated by microscopy 2 to 6 hours post-treatment.

### Overexpression of SOX9 in human epidermal melanocytes

Human epidermal melanocytes were transfected with either a human SOX9–expressing episomal plasmid or an empty vector control. The SOX9 expression vector was a customized pEBMulti-Puro episomal plasmid (FUJIFILM Wako Pure Chemical Corporation) carrying the human *SOX9* gene. Transfections were performed using the Neon NxT Electroporation System (#N1096, Thermo Fisher Scientific) in accordance with the manufacturer’s instructions. Each transfection reaction contained 2 µg of episomal vector DNA mixed with Lipofectamine LTX transfection reagent (#15338030, Thermo Fisher Scientific). At 24, 48, and 72 hours post-transfection, cells were fixed for immunofluorescence analysis to assess SOX9 overexpression. Fixed cells were incubated with primary rabbit antibodies against SOX9 (rabbit anti-SOX9, #AB5535, Sigma-Aldrich), Melan-A (mouse IgG2b anti-Melan-A, #917902, BioLegend, San Diego, CA, USA), and the His-tag (rabbit IgG1 anti-His-tag, #12698, Cell Signaling Technology, Danvers, MA, USA). The stained samples were then examined by fluorescence microscopy. The presence of robust nuclear SOX9 immunoreactivity and positive His-tag staining indicated successful overexpression of the *SOX9* transgene in the transfected melanocytes.

### *Ex vivo* culture of human skin explants

Full-thickness human skin explants were obtained from healthy donors and patients with vitiligo after obtaining informed consent. For experiments involving healthy skin (Fig. 4), tissues were collected from three independent donors. Each donor’s tissue was divided into multiple small explants, and each drug-treatment condition was assessed with three biological replicates (n = 3 per condition). For vitiligo skin (Fig. 6), a total of four 4-mm punch biopsies (two biopsies per patient) were obtained from the edge of lesional areas from two patients. Each biopsy was bisected, generating paired control and drug-treated samples, resulting in four biological replicates for each treatment condition (n = 2 per condition). Explants from healthy donors and patients with vitiligo were cultured and analyzed separately. Each skin explant was placed on a cell culture insert (8-μm pore size, #141082, Thermo Fisher Scientific) with the dermal side in contact with the culture medium and the epidermal side exposed to air. The culture medium consisted of DMEM (#044-29765, FUJIFILM Wako Pure Chemical Corporation) supplemented with 10% fetal bovine serum (#10270106, Thermo Fisher Scientific) and 1× penicillin-streptomycin solution (#168-23191, FUJIFILM Wako Pure Chemical Corporation). Explants were maintained at 37 °C in a humidified 5% CO_2_ incubator for 7 days, a duration well within the viability window reported for *ex vivo* human skin cultures. The culture medium was refreshed every 2 days. Drug treatments were administered by adding test compounds directly to the culture medium. Treatments were applied at the time of medium change to ensure consistent exposure, and the vehicle control consisted of an equivalent volume of DMSO in the medium.

### Drug treatment methods

For *in vitro* experiments, cells were cultured in Medium 254 (#M254500, Thermo Fisher Scientific)and treated for 24 to 48 hours with various compounds dissolved in DMSO. Control cells received an equivalent volume of DMSO as a vehicle control. Prior to these treatments, each compound underwent dose-response assays using serial dilutions to determine the highest concentration that was both nontoxic and effective. The optimal concentrations identified were used in all subsequent experiments and are detailed in Supplementary Table 6.

For *in vivo* experiments, compounds were dissolved in a 50:50 (v/v) ethanol/sesame oil mixture and topically applied daily to the dorsal skin of K14-Kitl mice for 2 weeks. Control mice received the vehicle solution without compound. Dose-response assays were conducted beforehand to establish the highest nontoxic and effective concentrations, which were then used in all experiments. The final working concentrations are listed in Supplementary Table 7.

For *ex vivo* experiments, skin explants were maintained for 1 week in DMEM supplemented with 10% fetal bovine serum and 1% penicillin-streptomycin. The culture medium, containing the respective compounds or DMSO vehicle, was replaced every 2 days. Prior dose-response assays determined the optimal nontoxic and effective concentrations for each compound, which were subsequently used in all experiments. These concentrations are provided in Supplementary Table 8.

### Statistical analysis

All experiments were repeated at least three times, and results are presented as the mean ± standard deviation. Statistical analyses were performed using unpaired two-tailed Student’s t-tests, one-way analysis of variance (ANOVA) with Dunnett’s *post hoc* test, or two-way ANOVA, as appropriate. The same statistical approach was applied to both *in vitro* and *in vivo* data. Statistical significance was defined as *P* < 0.05. All analyses were carried out using GraphPad Prism v9.5.3 (GraphPad Software, Boston, MA, USA).

### Data availability

The data that support the findings of this study are available from the corresponding author upon reasonable request.

## Acknowledgments

We gratefully acknowledge Hideki Nakagawa of the Research Support Platform at Osaka Metropolitan University Graduate School of Medicine for providing expert technical assistance with electron microscopy.

## Supplementary Information

### Extended Data Figures

**Extended Data Figure 1.**
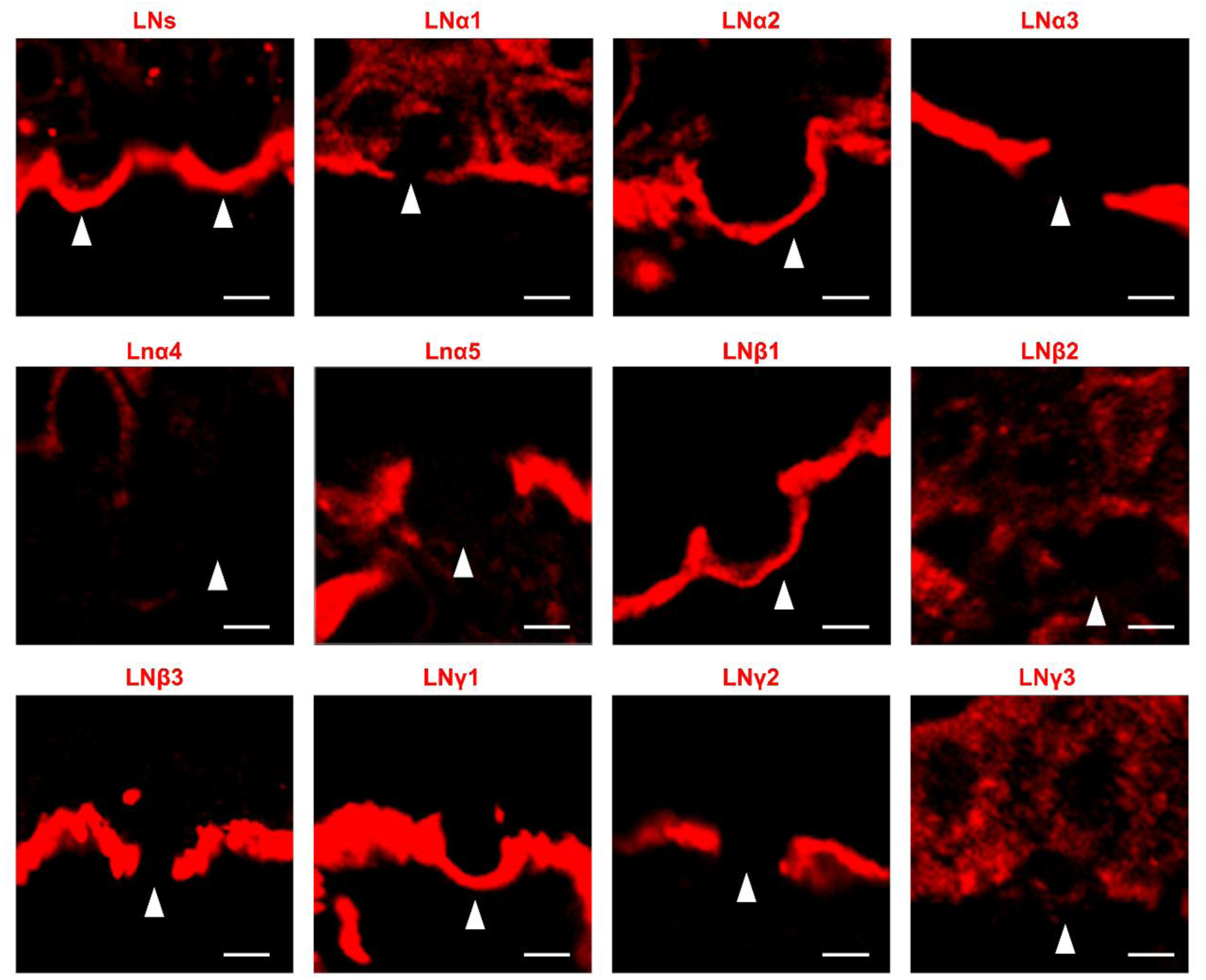
Immunofluorescent staining of various laminin chains at the dermal-epidermal junction in normal human skin. Antibodies were used to detect pan-laminins (LNs), laminin α chains (LNα1-LNα5), laminin β chains (LNβ1-LNβ3), and laminin γ chains (LNγ1-LNγ3). Melanocytes were identified using a melanocyte-specific marker and are indicated by white arrowheads. Each panel displays the localization pattern of an individual laminin chain. Scale bar: 5 μm.

**Extended Data Figure 2.**
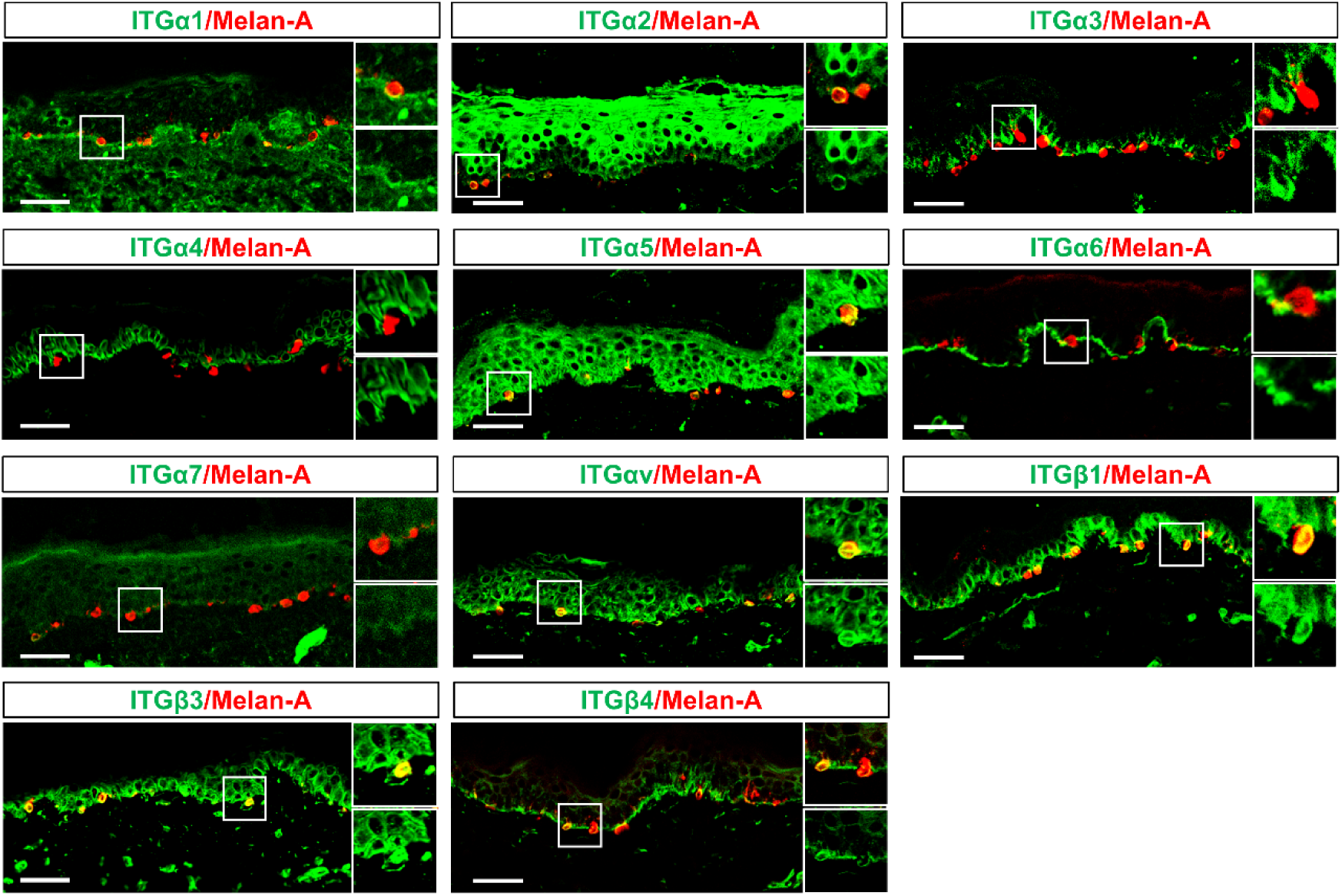
Expression patterns of integrin subunits in normal human melanocytes. Immunofluorescence staining of skin sections from three healthy controls showing the expression of various integrin subunits (green) in relation to melanocytes (Melan-A, red). Each panel represents a different integrin subunit: ITGα1, ITGα2, ITGα3, ITGα4, ITGα5, ITGα6, ITGα7, ITGαv, ITGβ1, ITGβ3, and ITGβ4. Insets show magnified views of the boxed regions, highlighting integrin localization in or around melanocytes at the dermal-epidermal junction. Scale bar: 50 μm.

**Extended Data Figure 3.**
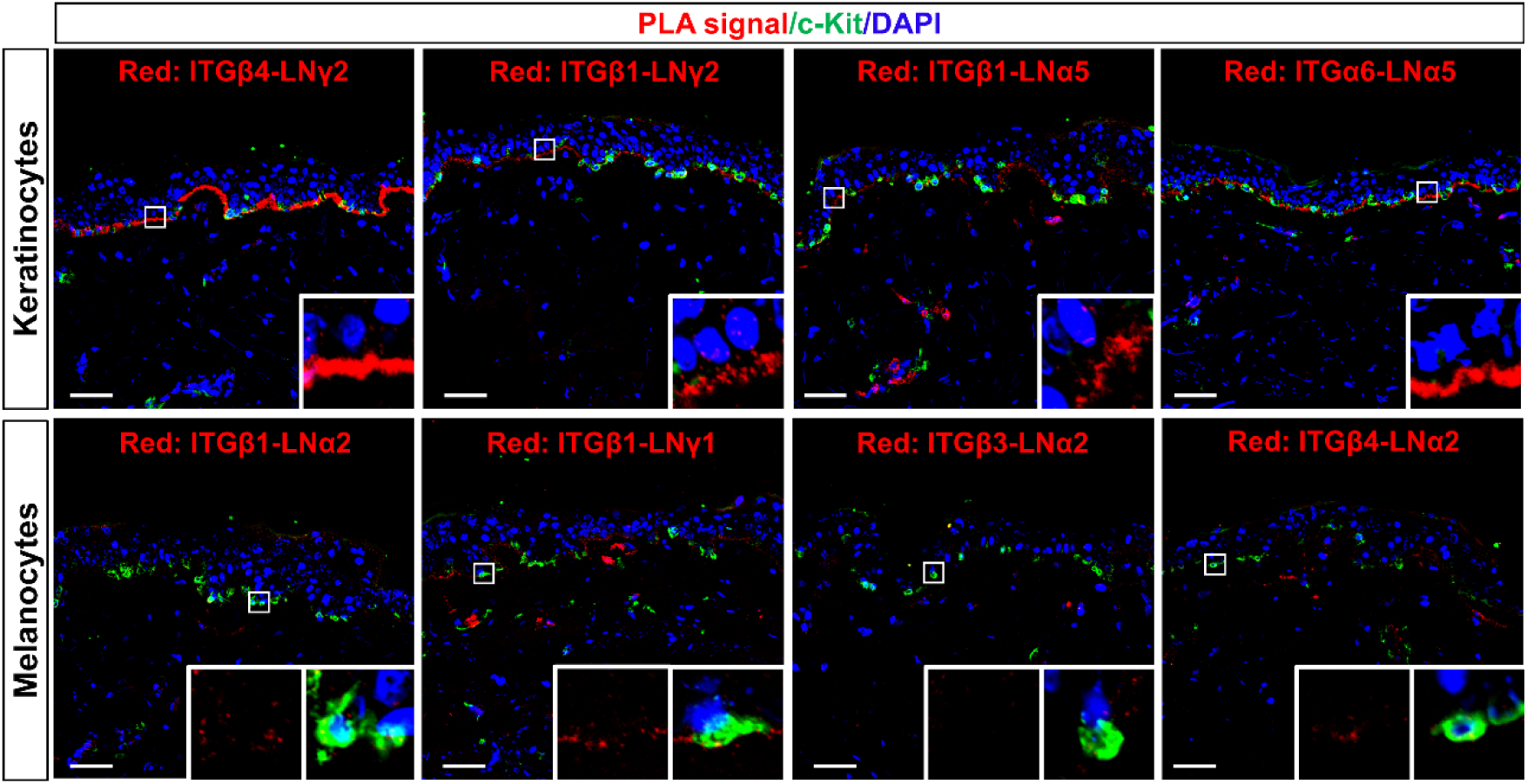
Analysis of integrin-laminin interactions in melanocytes from healthy human skin. Proximity ligation assay (PLA) detected protein-protein interactions between integrin subunits and laminin chains in skin sections from three healthy controls. Red signals represent PLA-positive puncta for specific integrin–laminin interactions. Upper panels show positive controls in keratinocytes, for ITGβ4-LNγ2 (a hemidesmosomal component), ITGβ1-LNγ2, ITGβ1-LNα5, and ITGα6-LNα5. Lower panels show PLA signals for integrin–laminin α2 and integrin–laminin γ1 interactions in melanocytes, identified by c-Kit (green). Nuclei are counterstained with 4′,6-diamidino-2-phenylindole (DAPI; blue). Insets show magnified views of boxed areas. Scale bar: 50 μm.

**Extended Data Figure 4.**
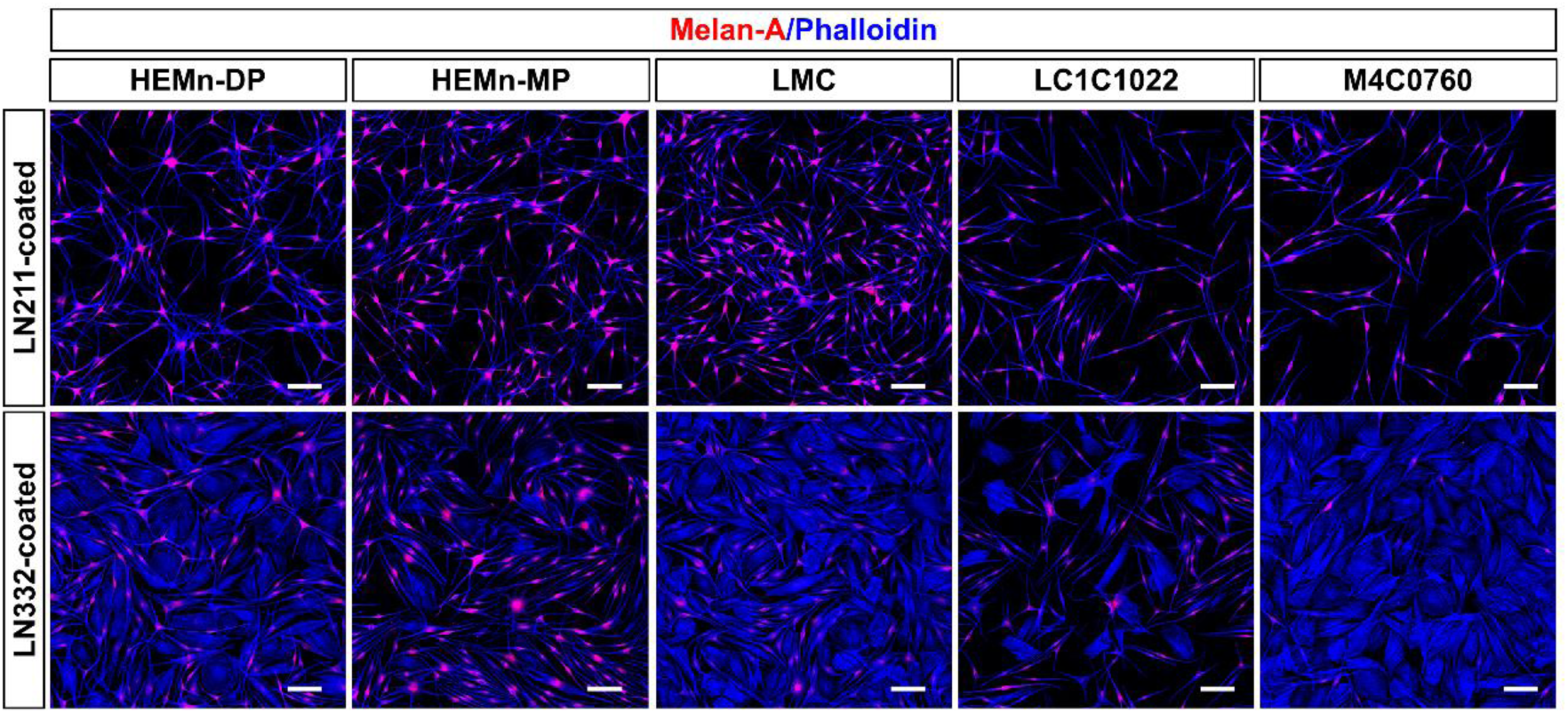
Laminin-332 (LN332) induces morphological changes and alters melanogenic protein expression in multiple human melanocyte lines. Representative immunofluorescence images of five melanocyte lines (HEMn-DP, HEMn-MP, LMC, LC1C1022, and M4C0760) cultured on plates coated with either laminin-211 (LN211) or LN332. Cells were stained for Melan-A (red) to assess melanocyte identity and melanogenic status and for phalloidin (blue) to visualize actin cytoskeleton and overall cell morphology following LN332 treatment. Scale bar: 100 μm.

**Extended Data Figure 5.**
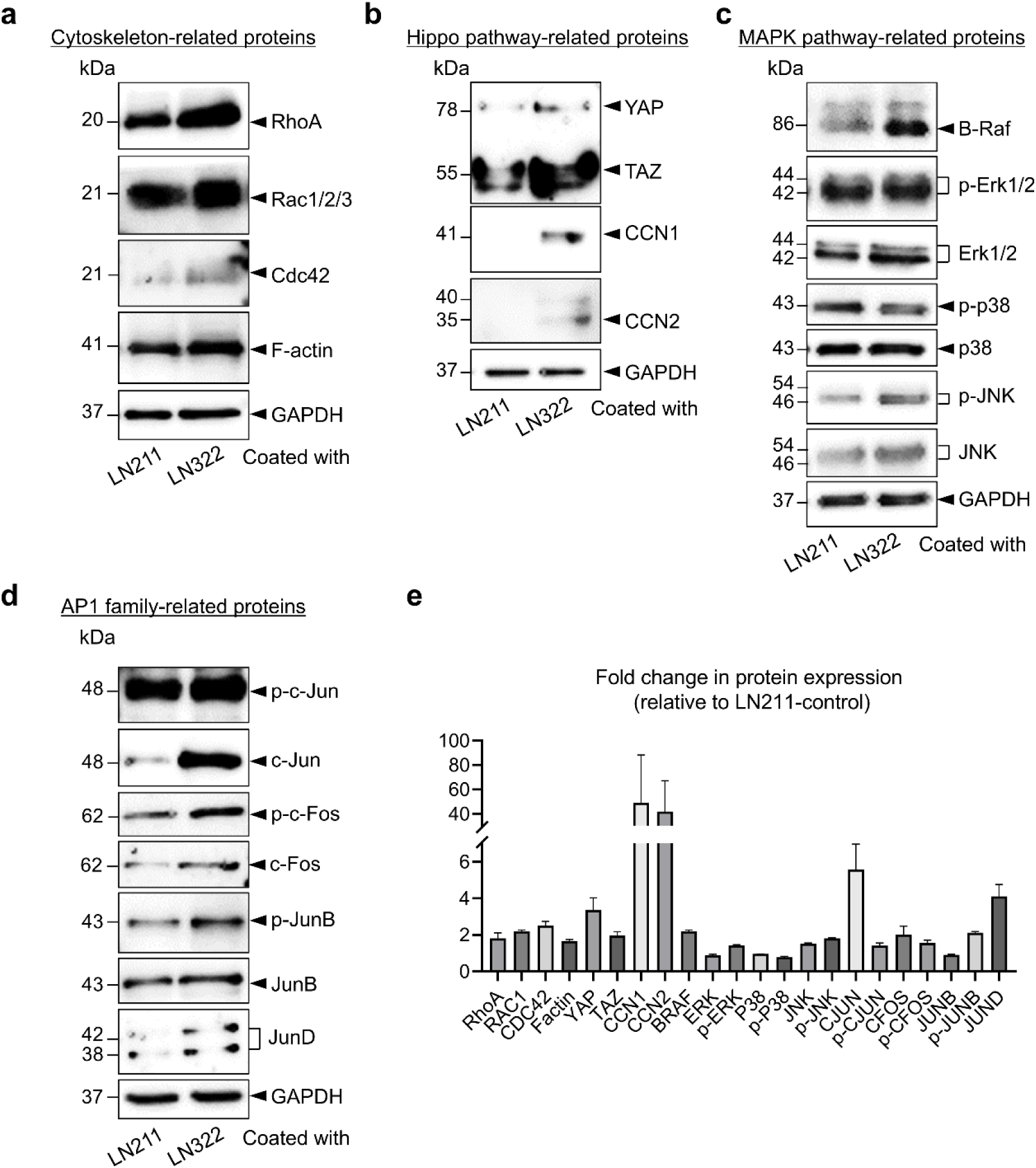
Western blot analysis of cytoskeleton components, Hippo pathway, MAPK pathway, and AP-1 family members in laminin-332 (LN332)-induced dedifferentiated melanocytes. **a-d.** Representative immunoblots showing expression levels of proteins related to the cytoskeleton (**a**), Hippo pathway (**b**), MAPK pathway (**c**), and AP-1 family factors (**d**). Lane 1: melanocytes cultured on laminin-211 (LN211; control). Lane 2: melanocytes cultured on LN332. Glyceraldehyde 3-phosphate dehydrogenase (GAPDH) was used as a loading control. **e.** Quantification of protein expression levels relative to GAPDH and normalized to LN211 control. Data are presented as mean ± SEM from three independent experiments.

**Extended Data Figure 6.**
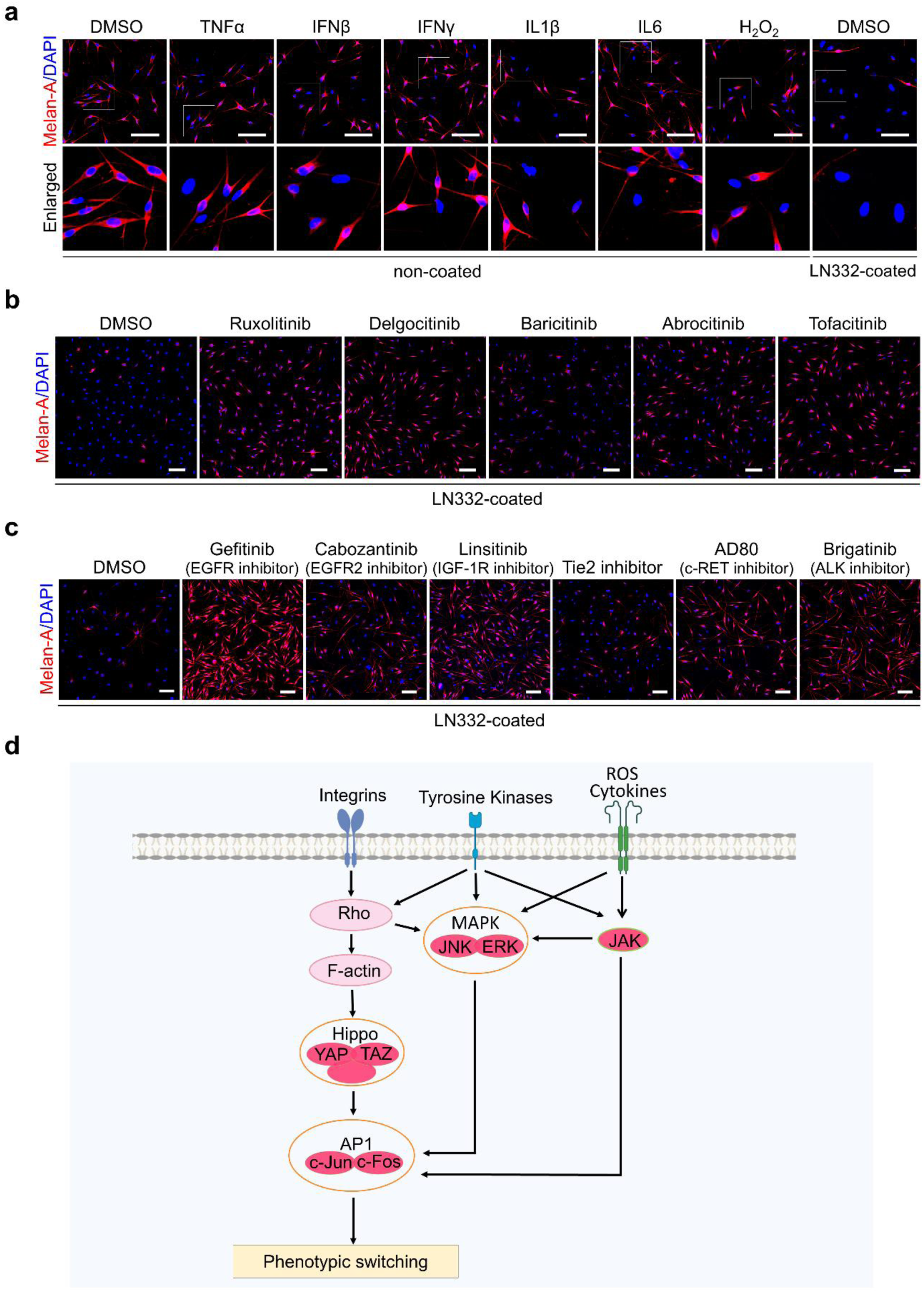
Influence of vitiligo-associated factors and pharmacological inhibitors on melanocyte dedifferentiation. **a.** Immunofluorescence analysis of Melan-A (red) expression in melanocytes treated with vitiligo-associated stressors, including proinflammatory cytokines (TNFα, IFNβ, IFNγ, IL1β, IL6) and oxidative stress (H_2_O_2_). Melanocytes were cultured on either noncoated surfaces (left panels) or surfaces coated with laminin-332 (LN332; right panels). Lower panels show enlarged views of selected regions. Scale bar: 100 μm. DAPI, 4′,6-diamidino-2-phenylindole; DMSO, dimethyl sulfoxide. **b.** Effects of JAK inhibitors (ruxolitinib, delgocitinib, baricitinib, abrocitinib, and tofacitinib) on LN332-induced melanocyte dedifferentiation. Melan-A (red) expression was used to assess melanocyte differentiation status. Scale bar: 100 μm. **c.** Effects of tyrosine kinase inhibitors targeting EGFR, EGFR2, IGF-1R, Tie2, c-RET, and ALK on LN332-induced dedifferentiation. Scale bar: 100 μm. **d.** Schematic diagram summarizing the signaling pathways implicated in melanocyte dedifferentiation. ROS, reactive oxygen species.

**Extended Data Figure 7.**
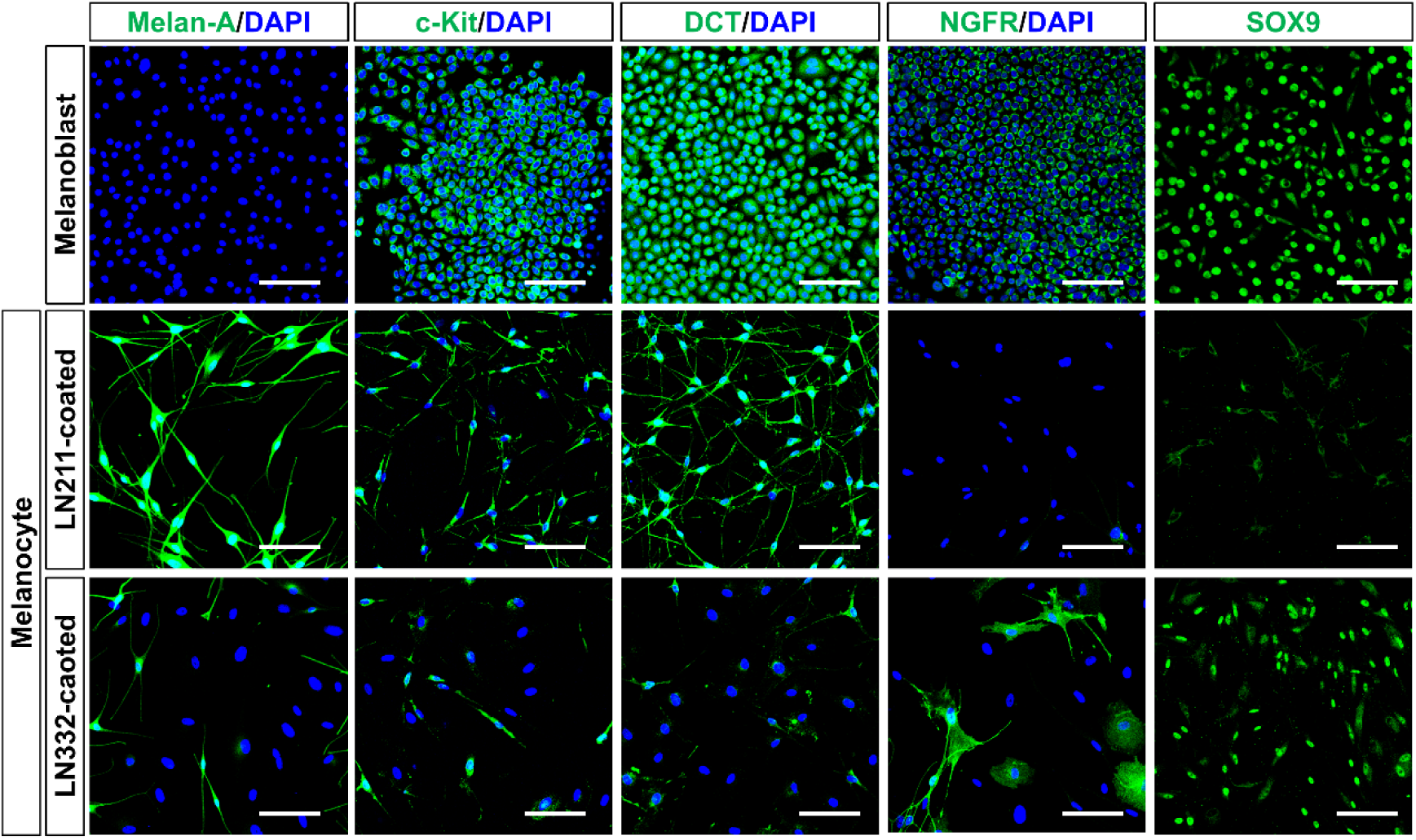
Comparative expression of precursor and melanocyte markers in melanoblasts and melanocytes cultured on laminin-211 (LN211) or laminin-332 (LN332) substrates. Immunofluorescence staining of melanoblasts (top row) and differentiated melanocytes cultured on plates coated with LN211 (middle row) or LN332 (bottom row). Cells were stained for Melan-A, c-Kit, DCT, NGFR, and SOX9 (green) to evaluate the expression of melanocyte lineage and precursor markers. Nuclei were counterstained with 4′,6-diamidino-2-phenylindole (DAPI; blue). Scale bar: 100 μm.

**Extended Data Figure 8.**
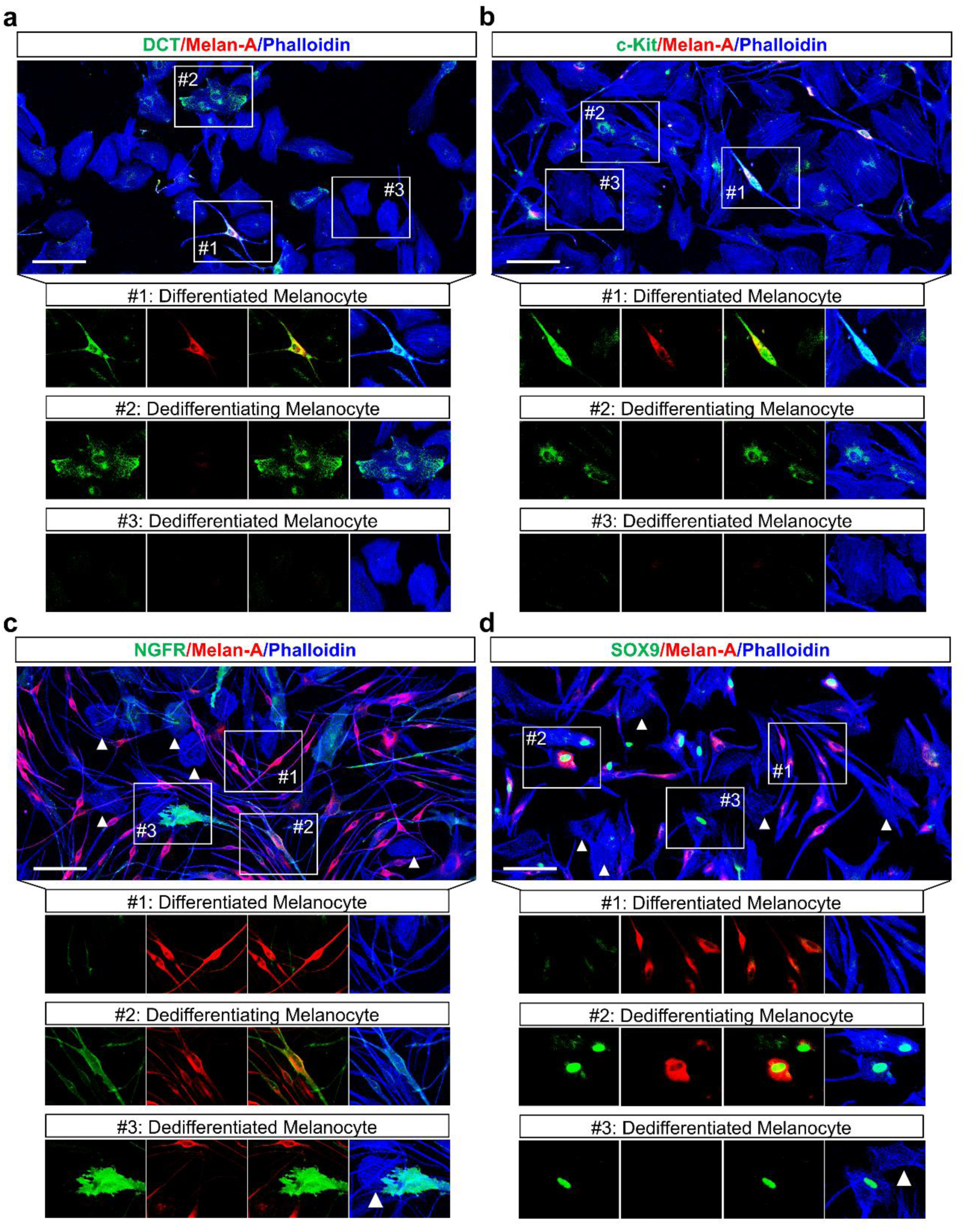
Multiparameter immunostaining reveals distinct melanocyte subpopulations following laminin-332 (LN332) treatment. Representative immunofluorescence images of melanocytes cultured on LN332-coated plates and stained with combinations of melanocyte lineage and dedifferentiation markers: **a.** DCT (green), Melan-A (red), and phalloidin (blue); **b.** c-Kit (green), Melan-A (red), and phalloidin (blue); **c.** NGFR (green), Melan-A (red), and phalloidin (blue); **d.** SOX9 (green), Melan-A (red), and phalloidin (blue). Three representative melanocyte states are shown in magnified insets below each panel. #1: Differentiated melanocytes; #2: dedifferentiating melanocytes; #3: dedifferentiated melanocytes. Arrowheads in **c** indicate NGFR/Melan-A double-negative dedifferentiated cells. Arrowheads in **d** indicate SOX9/Melan-A double-negative dedifferentiated cells. Scale bar: 100 μm.

**Extended Data Figure 9.**
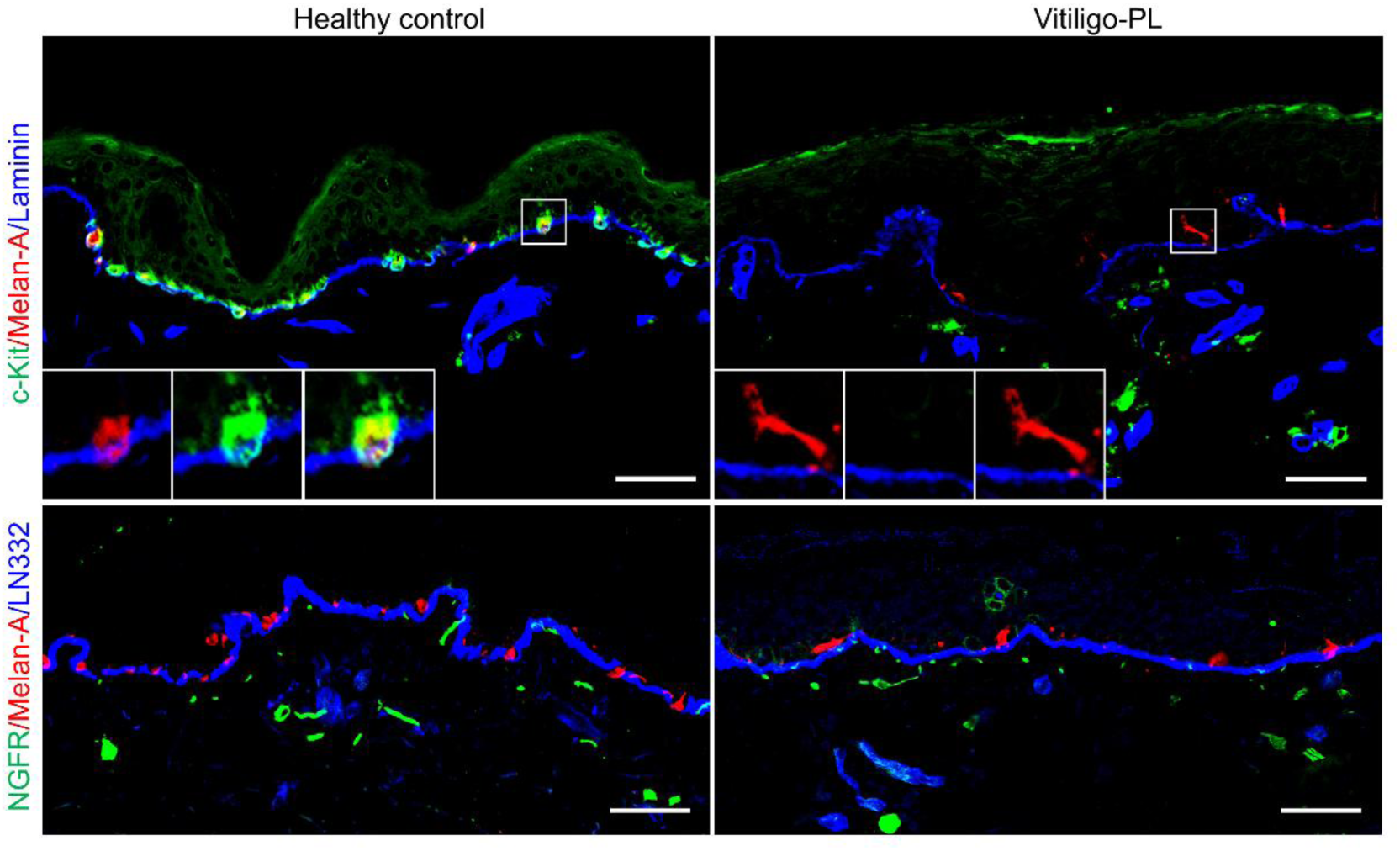
Identification of dedifferentiating and dedifferentiated melanocytes in vitiligo skin. Immunofluorescence staining of c-Kit (green), NGFR (green), Melan-A (red), and laminin or laminin-332 (LN332; blue) in healthy control and perilesional vitiligo (Vitiligo-PL) skin. Insets highlight melanocytes. Scale bar: 50 μm.

**Extended Data Figure 10.**
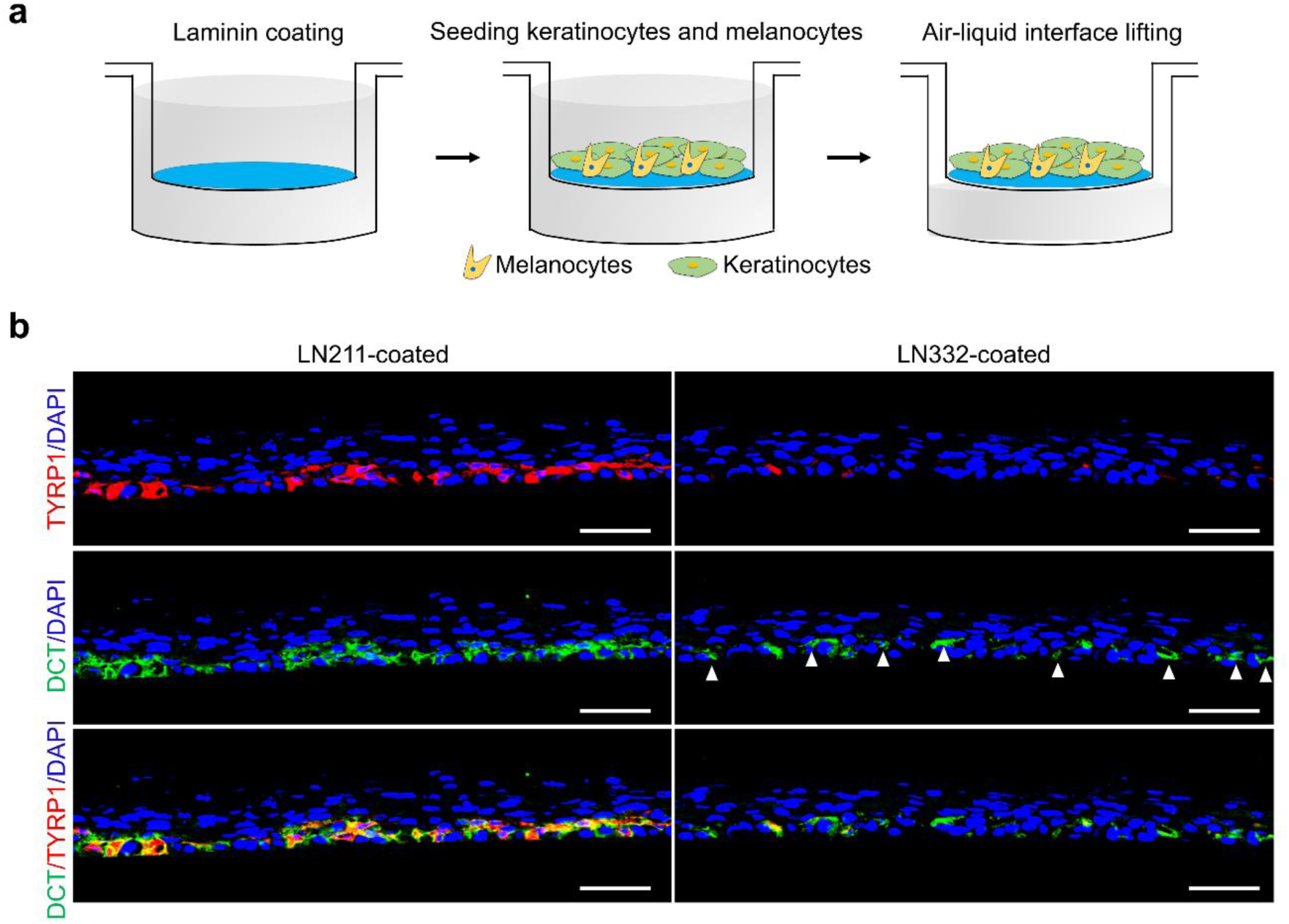
Laminin-332 (LN332) reduces mature melanocytes and increases dedifferentiating melanocytes in a three-dimensional (3D) skin model. **a.** Schematic overview of the protocol for constructing a 3D reconstructed human skin model. Keratinocytes and melanocytes were coseeded onto laminin-coated transwell inserts, followed by culture at the air-liquid interface to promote epidermal stratification. **b.** Immunofluorescence staining of TYRP1 (red), DCT (green), and 4′,6-diamidino-2-phenylindole (DAPI; blue) in 3D skin models cultured on substrates coated with laminin-211 (LN211) or LN332. Arrowheads indicate DCT⁺/TYRP1⁻ melanocytes observed in the LN332-coated model, representing dedifferentiating cells. Scale bar: 50 μm.

**Extended Data Figure 11.**
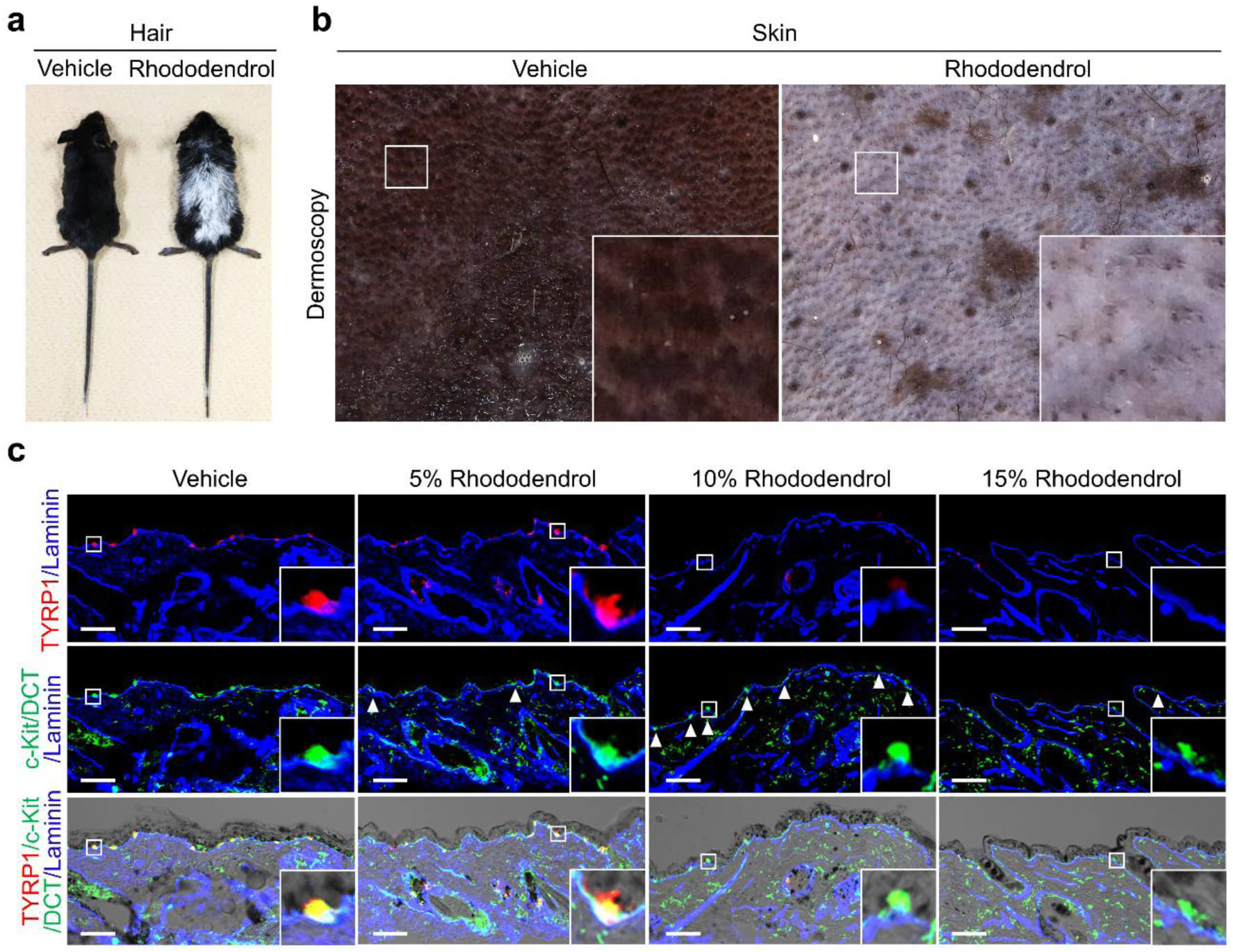
Dynamics of melanocyte dedifferentiation in a rhododendrol-induced leukoderma mouse model. **a.** Macroscopic appearance of K14-Kitl mice after topical application of vehicle or rhododendrol. Mice treated with rhododendrol exhibit pronounced hair depigmentation. **b.** Dermoscopic images of dorsal skin from mice treated with vehicle or rhododendrol, showing depigmented patches in rhododendrol-treated skin. **c.** Immunofluorescence staining of TYRP1 (red), DCT (green), c-Kit (green), and laminin (blue) in mouse dorsal skin treated with increasing concentrations of rhododendrol (5%, 10%, 15%). Scale bar: 50 μm. Arrowheads indicate c-Kit^+^/TYRP1^−^ or DCT^+^/TYRP1^−^ dedifferentiating melanocytes.

**Extended Data Figure 12.**
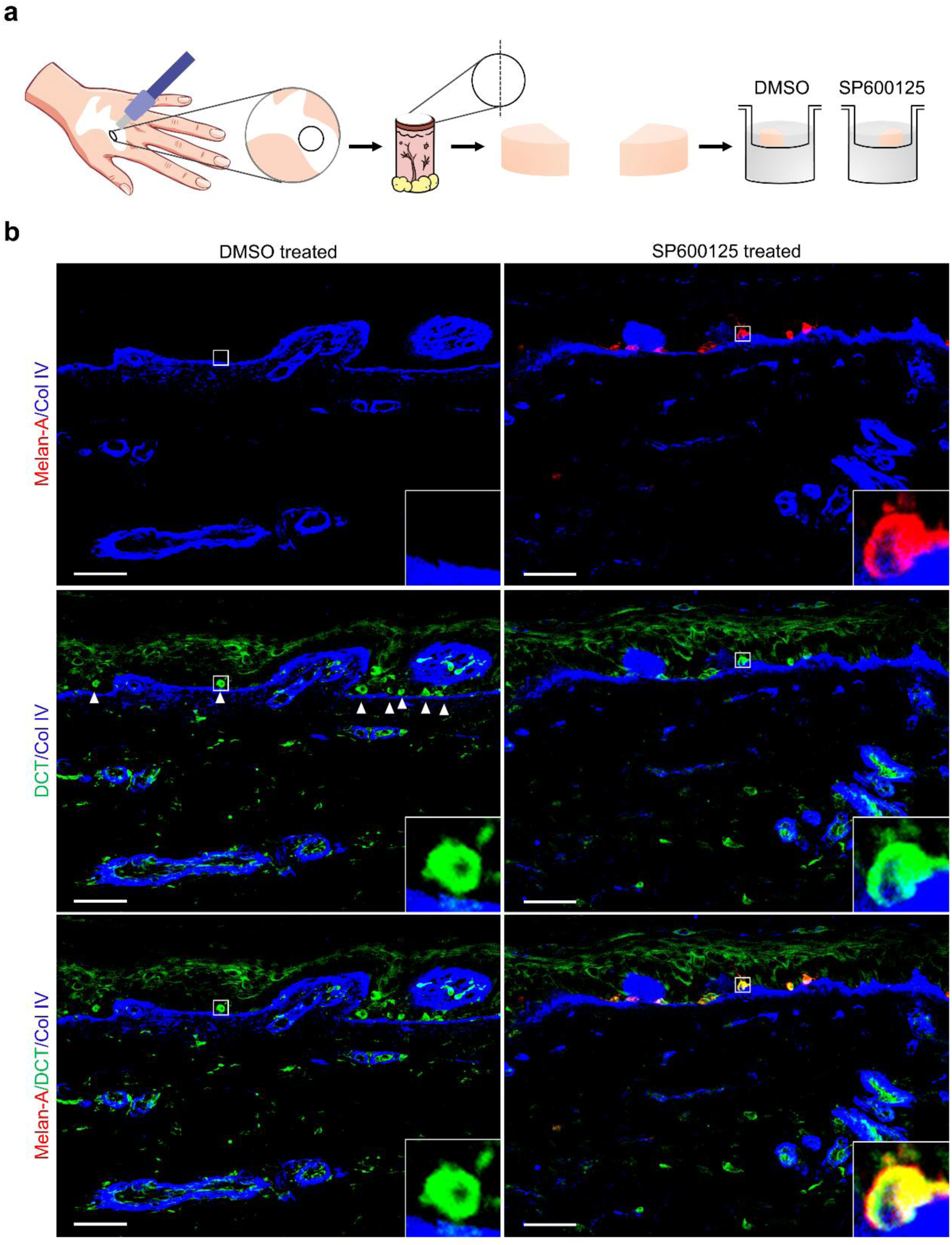
SP600125 promotes redifferentiation of melanocytes in *ex vivo*-cultured skin from patients with vitiligo. **a.** Schematic illustration of the *ex vivo* culture system using lesional skin from patients with vitiligo. Skin explants were sectioned and cultured for 1 week in the presence of dimethyl sulfoxide (DMSO; vehicle control) or the JNK inhibitor SP600125. **b.** Immunofluorescence staining of Melan-A (red), DCT (green), and type IV collagen (Col IV, blue) in *ex vivo*-cultured vitiligo skin. Arrowheads indicate DCT⁺/Melan-A⁻ dedifferentiating melanocytes, which are prevalent in DMSO-treated epidermis. Inset images show higher magnification of boxed regions, illustrating representative dedifferentiated and redifferentiated melanocytes. Scale bar: 50 μm.

**Extended Data Figure 13.**
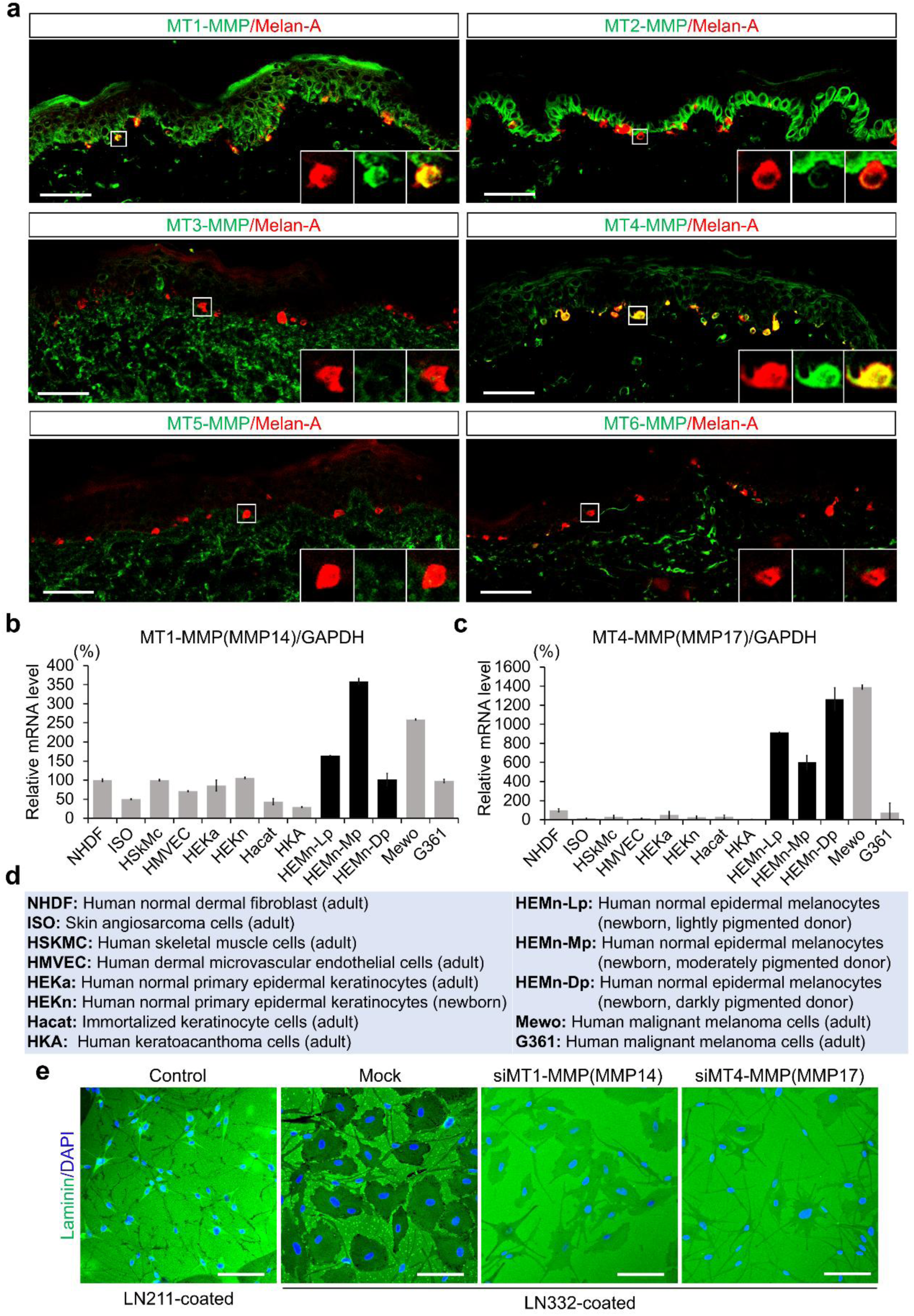
Analysis of membrane-type matrix metalloproteinase (MT-MMP)-mediated laminin-332 (LN332) degradation by melanocytes. **a.** Immunofluorescence staining of MT1-MT6-MMPs (green) and Melan-A (red) in normal human epidermis. Insets show representative melanocytes coexpressing specific MT-MMPs. Scale bar: 50 μm. **b-c.** Quantitative PCR analysis of MT1-MMP (MMP14) (**b**) and MT4-MMP (MMP17) (**c**) mRNA expression across various human cell types. GAPDH, glyceraldehyde 3-phosphate dehydrogenase. **d.** Cell line abbreviations used in **b** and **c**. **e.** LN332 degradation assay using control and MT-MMP-silenced melanocytes. Cells were cultured on LN332-coated plates and stained for laminin (green) and nuclei (4′,6-diamidino-2-phenylindole, DAPI; blue). Scale bar: 100 μm.

**Extended Data Figure 14.**
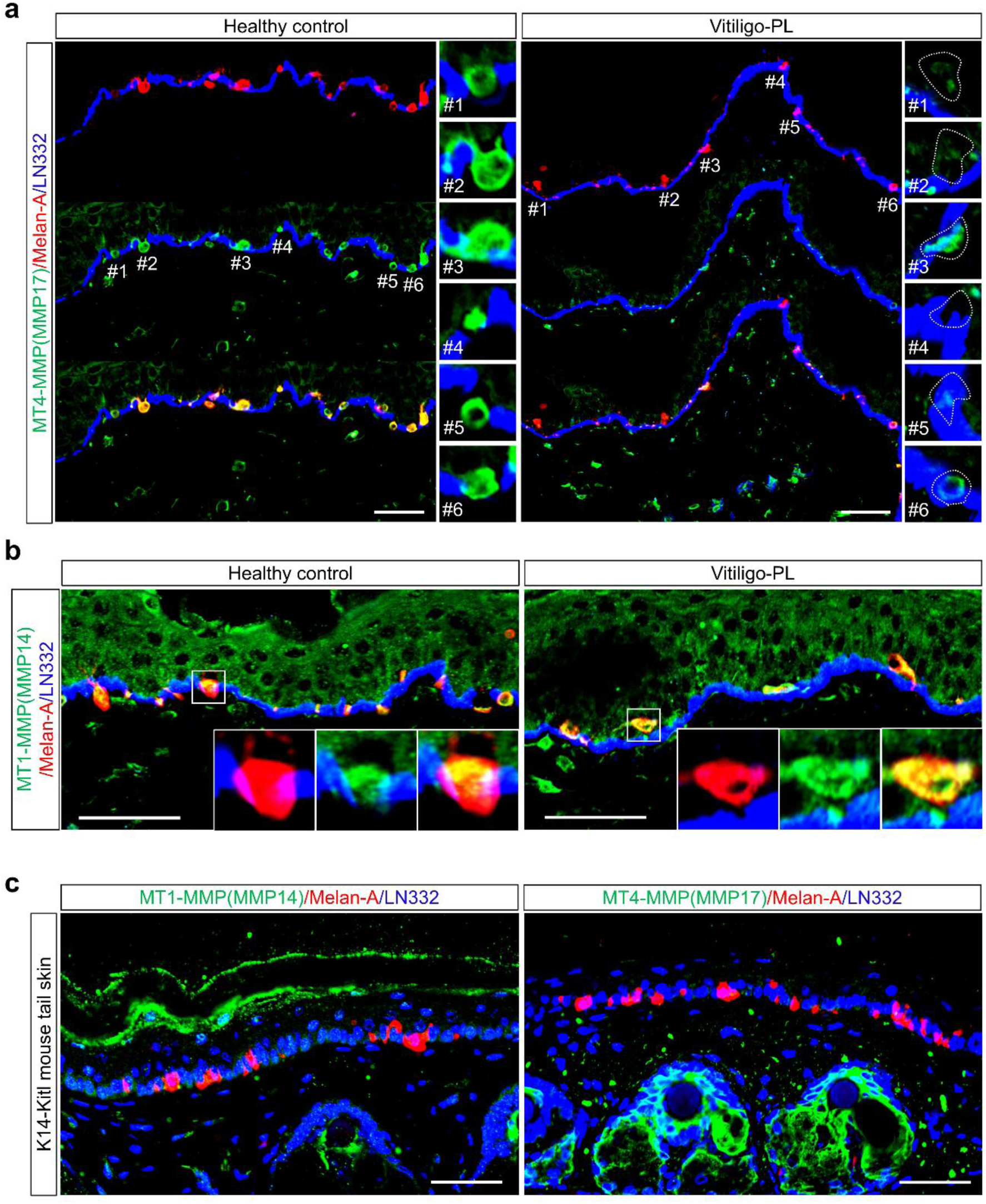
Analysis of membrane-type matrix metalloproteinase (MT-MMP) expression in melanocytes from the skin of patients with vitiligo and from mice. **a.** Immunofluorescence staining of MT4-MMP (MMP17), Melan-A, and laminin-332 (LN332) in healthy control and perilesional vitiligo (Vitiligo-PL) human skin. Insets on the right show representative melanocytes (#1-6). Scale bar: 50 μm. **b.** Immunofluorescence staining of MT1-MMP (MMP14), Melan-A, and LN332 in healthy control and Vitiligo-PL human skin. Insets show representative melanocytes. Scale bar: 50 μm. **c.** Immunostaining of MT1-MMP (left) or MT4-MMP (right), Melan-A, and LN332 in K14-Kitl mouse tail skin to assess MT-MMP expression *in vivo*. Scale bar: 50 μm.

**Extended Data Figure 15.**
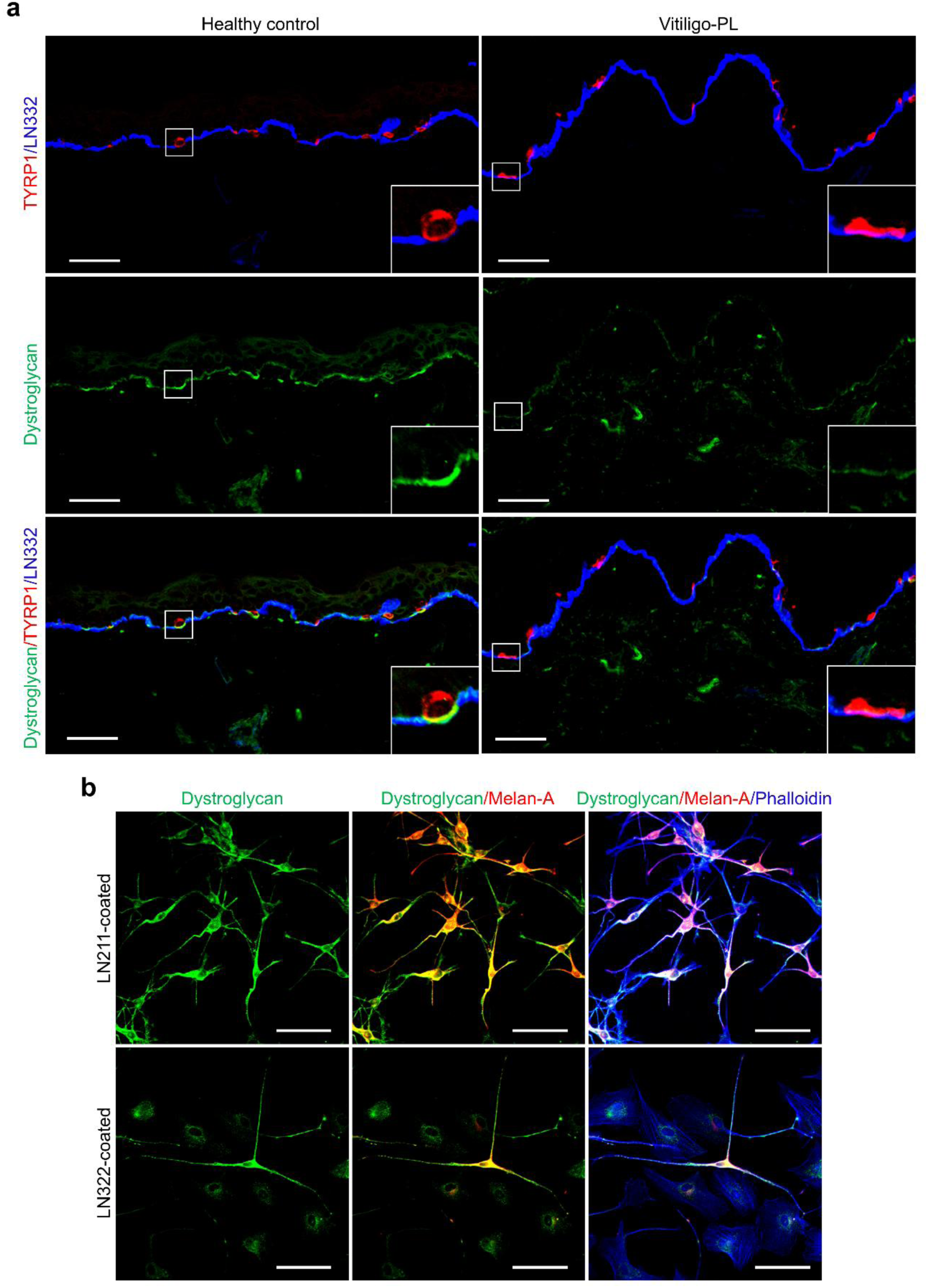
Analysis of dystroglycan expression in vitiligo skin and cultured melanocytes. **a.** Immunofluorescence staining of dystroglycan (green), TYRP1 (red), and laminin-332 (LN332; blue) in healthy control and perilesional vitiligo (Vitiligo-PL) skin. Insets enlarge melanocytes. Scale bar: 50 μm. **b.** Expression of dystroglycan in melanocytes cultured on substrates coated with laminin-211 (LN211) or LN332. Immunostaining was performed for dystroglycan (green), Melan-A (red), and phalloidin (blue). Scale bar: 100 μm.

**Extended Data Figure 16.**
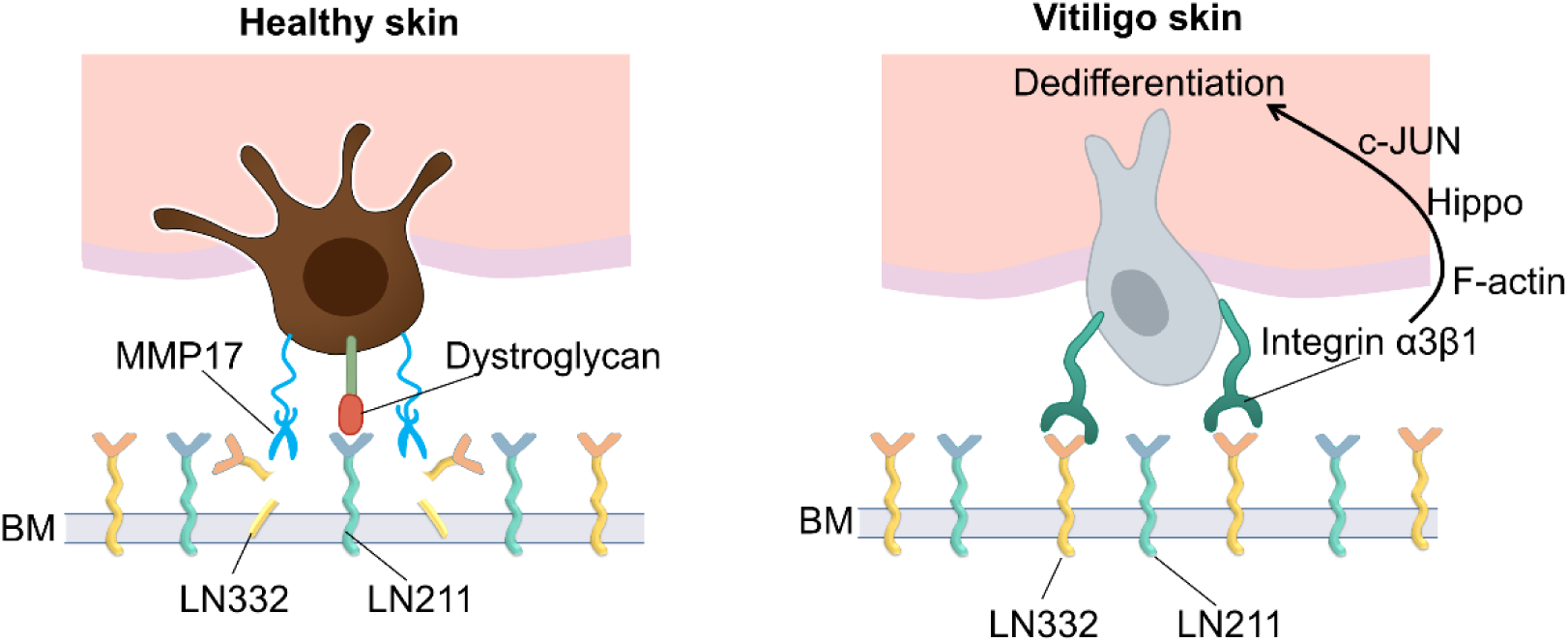
Schematic illustration of melanocyte dedifferentiation induced by the vitiligo-associated microenvironment. In healthy skin (left), melanocytes adhere to the basement membrane (BM) through dystroglycan-laminin-211 interactions and maintain their differentiated state. MMP17 expression contributes to localized remodeling without disrupting homeostasis. In vitiligo skin (right), increased laminin-332 (LN332) exposure and integrin α3β1 engagement activate intracellular signaling cascades involving F-actin reorganization and the Hippo and MAPK pathways, ultimately converging on c-Jun activation. This signaling switch promotes melanocyte dedifferentiation.

## Supplementary Tables

**Table S1.**
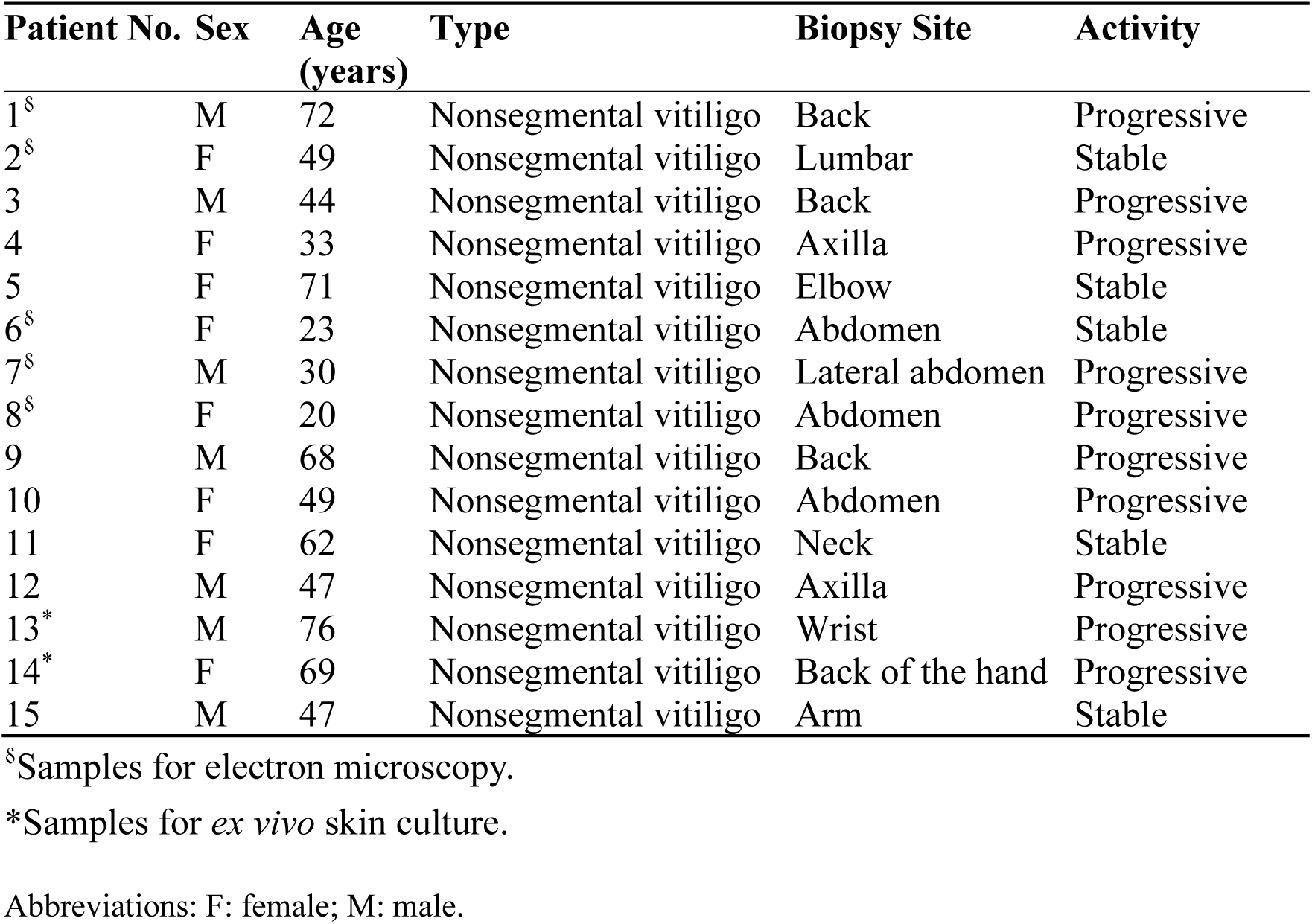
Clinical characteristics of the patients.

| Patient No. | Sex | Age<br>(years) | Type | Biopsy Site | Activity |
| --- | --- | --- | --- | --- | --- |
| 1<sup>§</sup> | M | 72 | Nonsegmental vitiligo | Back | Progressive |
| 2<sup>§</sup> | F | 49 | Nonsegmental vitiligo | Lumbar | Stable |
| 3 | M | 44 | Nonsegmental vitiligo | Back | Progressive |
| 4 | F | 33 | Nonsegmental vitiligo | Axilla | Progressive |
| 5 | F | 71 | Nonsegmental vitiligo | Elbow | Stable |
| 6<sup>§</sup> | F | 23 | Nonsegmental vitiligo | Abdomen | Stable |
| 7<sup>§</sup> | M | 30 | Nonsegmental vitiligo | Lateral abdomen | Progressive |
| 8<sup>§</sup> | F | 20 | Nonsegmental vitiligo | Abdomen | Progressive |
| 9 | M | 68 | Nonsegmental vitiligo | Back | Progressive |
| 10 | F | 49 | Nonsegmental vitiligo | Abdomen | Progressive |
| 11 | F | 62 | Nonsegmental vitiligo | Neck | Stable |
| 12 | M | 47 | Nonsegmental vitiligo | Axilla | Progressive |
| 13<sup>*</sup> | M | 76 | Nonsegmental vitiligo | Wrist | Progressive |
| 14<sup>*</sup> | F | 69 | Nonsegmental vitiligo | Back of the hand | Progressive |
| 15 | M | 47 | Nonsegmental vitiligo | Arm | Stable |

<sup>§</sup>Samples for electron microscopy.
<sup>\*</sup>Samples for *ex vivo* skin culture.
Abbreviations: F: female; M: male.

**Table S2.** List of primary antibodies used for immunofluorescence and immunohistochemical analyses.

| Antibody Name | Host, Isotype, and Subclass | Dilution | Catalog No. | Company Name |
| --- | --- | --- | --- | --- |
| Laminin | Rabbit | 1:100 | ab11575 | Abcam |
| Laminin α1 | Mouse IgG1 | 1:100 | AMAb91091 | Sigma-Aldrich |
| Laminin α2 | Rat IgG1 | 1:100 | L0663 | Sigma-Aldrich |
| Laminin α3 | Mouse IgG2a | 1:100 | FDV-0024 | Funakoshi |
| Laminin α4 | Rabbit | 1:100 | SAB4501719 | Sigma-Aldrich |
| Laminin α5 | Mouse IgG1 | 1:100 | MABT39 | Sigma-Aldrich |
| Laminin β1 | Rat IgG1 | 1:100 | ab44941 | Abcam |
| Laminin β2 | Mouse IgG1 | 1:100 | AMAb91096 | Sigma-Aldrich |
| Laminin β3 | Rabbit | 1:100 | 26795-1-AP | Proteintech |
| Laminin γ1 | Rat IgG2a | 1:100 | ab80580 | Abcam |
| Laminin γ2 | Rabbit | 1:100 | SAB2700481 | Sigma-Aldrich |
| Laminin-332 | Mouse IgG1 | 1:100 | ab78286 | Abcam |
| Integrin α1 | Mouse IgG1 | 1:100 | NBP2-76478 | Novus Biologicals |
| Integrin α2 | Mouse IgG1 | 1:100 | NBP2-76483 | Novus Biologicals |
| Integrin α3 | Mouse IgG2a | 1:100 | NBP1-97691 | Novus Biologicals |
| Integrin α4 | Rabbit | 1:100 | 19676-1-AP | Proteintech |
| Integrin α5 | Rabbit | 1:100 | 27224-1-AP | Proteintech |
| Integrin α6 | Mouse IgG2b | 1:100 | ab20142 | Abcam |
| Integrin α7 | Mouse IgG1 | 1:100 | ab195959 | Abcam |
| Integrin αv | Rabbit | 1:100 | 100563-T08 | Sino Biological |
| Integrin β1 | Mouse IgG2a | 1:100 | ab30388 | Abcam |
| Integrin β1 | Rabbit | 1:100 | PA532887 | Thermo Fisher |
| Integrin β3 | Rabbit | 1:100 | 100458-T08 | Sino Biological |
| Integrin β4 | Rabbit | 1:100 | PB9007 | Boster Bio |
| Dystroglycan | Rabbit | 1:100 | 11017-1-AP | Proteintech |
| DDR | Goat | 1:100 | AF2396 | R&D Systems |
| Melan-A | Mouse IgG2b | 1:100 | 917901 | BioLegend |
| c-Kit | Goat | 1:100 | AF332 | R&D Systems |
| c-Kit | Rat IgG2b | 1:100 | NBP1-43359 | Novus Biologicals |
| PMEL17 | Rabbit | 1:100 | ab137078 | Abcam |
| TYRP1 | Mouse IgG2a | 1:100 | 917801 | BioLegend |
| Neurofilament | Rabbit | 1:100 | 2837 | Cell Signaling Technology |
| COL2 | Rabbit | 1:100 | 28459-1-AP | BioLegend |
| α-SMA | Mouse IgG2a | 1:100 | M0851 | Dako |
| SOX9 | Rabbit | 1:100 | AB5535 | Sigma-Aldrich |
| NGFR | Rabbit | 1:100 | ab52987 | Abcam |
| DCT | Mouse IgG2a | 1:100 | sc-74439 | Santa Cruz Biotechnology |
| Collagen IV | Rabbit | 1:100 | ab19808 | Abcam |
| F-actin | Mouse IgM | 1:100 | ab205 | Abcam |
| p-JUN | Rabbit | 1:100 | 3270S | Cell Signaling Technology |
| His-Tag | Rabbit | 1:100 | 12698 | Cell Signaling Technology |
| MT1-MMP | Rabbit | 1:100 | ARG57921 | Arigo Biolaboratories |
| MT2-MMP | Rabbit | 1:100 | HPA040390 | Sigma-Aldrich |
| MT3-MMP | Rabbit | 1:100 | HPA023693 | Sigma-Aldrich |
| MT4-MMP | Rabbit | 1:100 | PA5-28219 | Thermo Fisher |
| MT5-MMP | Rabbit | 1:100 | AB924 | R&D Systems |
| MT6-MMP | Mouse IgG1 | 1:100 | MAB11421 | R&D Systems |
| Rabbit IgG Control | Rabbit | 1:100 | ab172730 | Abcam |
| Mouse IgG Control | Mouse | 1:100 | ab37355 | Abcam |
| Alexa Fluor 633 Phalloidin |  | 1:500 | A22284 | Thermo Fisher |

**Table S3.** List of secondary antibodies used for immunofluorescence and immunohistochemical analyses.

| **Antibody Name** | **Host** | **Dilution** | **Catalog No.** | **Company Name** |
| --- | --- | --- | --- | --- |
| Alexa Fluor Plus 488 Goat Anti-Rabbit IgG (H+L) | Goat | 1:1,000 | A32731 | Invitrogen |
| Alexa Fluor Plus 555 Goat Anti-Rabbit IgG (H+L) | Goat | 1:1,000 | A32732 | Invitrogen |
| Alexa Fluor Plus 647 Goat Anti-Rabbit IgG (H+L) | Goat | 1:1,000 | A32733 | Invitrogen |
| Alexa Fluor Plus 488 Goat Anti-Mouse IgG (H+L) | Goat | 1:1,000 | A32723 | Invitrogen |
| Alexa Fluor Plus 555 Goat Anti-Mouse IgG (H+L) | Goat | 1:1,000 | A32727 | Invitrogen |
| Alexa Fluor Plus 647 Goat Anti-Mouse IgG (H+L) | Goat | 1:1,000 | A21236 | Invitrogen |
| Goat Anti-Mouse IgG1, Alexa Fluor 488 | Goat | 1:1,000 | A21121 | Invitrogen |
| Goat Anti-Mouse IgG1, Alexa Fluor 555 | Goat | 1:1,000 | A21127 | Invitrogen |
| Goat Anti-Mouse IgG1, Alexa Fluor 647 | Goat | 1:1,000 | A21240 | Invitrogen |
| Goat Anti-Mouse IgG2a, Alexa Fluor 488 | Goat | 1:1,000 | A21131 | Invitrogen |
| Goat Anti-Mouse IgG2a, Alexa Fluor 555 | Goat | 1:1,000 | A21137 | Invitrogen |
| Goat Anti-Mouse IgG2a, Alexa Fluor 647 | Goat | 1:1,000 | A21136 | Invitrogen |
| Goat Anti-Mouse IgG2b, Alexa Fluor 488 | Goat | 1:1,000 | A21141 | Invitrogen |
| Goat Anti-Mouse IgG2b, Alexa Fluor 555 | Goat | 1:1,000 | A21147 | Invitrogen |
| Goat Anti-Mouse IgG2b, Alexa Fluor 647 | Goat | 1:1,000 | A21242 | Invitrogen |
| Goat Anti-Rat IgG (H+L), Alexa Fluor 488 | Goat | 1:1,000 | A11006 | Invitrogen |
| Goat Anti-Rat IgG (H+L), Alexa Fluor 555 | Goat | 1:1,000 | A48263 | Invitrogen |
| Goat Anti-Rat IgG (H+L), Alexa Fluor 647 | Goat | 1:1,000 | A21247 | Invitrogen |
| Donkey Anti-Goat IgG (H+L), Alexa Fluor Plus 488 | Donkey | 1:1,000 | A32814 | Invitrogen |
| Donkey Anti-Rabbit IgG (H+L), Alexa Fluor Plus 555 | Donkey | 1:1,000 | A32794 | Invitrogen |
| Donkey Anti-Mouse IgG (H+L), Alexa Fluor Plus 647 | Donkey | 1:1,000 | A32787 | Invitrogen |

**Table S4.** List of primary antibodies used for Western blot analyses.

| **Antibody Name** | **Host, Isotype, and Subclass** | **Dilution** | **Catalog No.** | **Company Name** |
| --- | --- | --- | --- | --- |
| c-Kit | Rabbit | 1:1,000 | 3074 | Cell Signaling Technology |
| TYR | Mouse IgG2a | 1:1,000 | SC20035 | Santa Cruz Biotechnology |
| Melan-A | Rabbit | 1:100 | ab51061 | Abcam |
| TYRP1 | Mouse IgG2a | 1:100 | sc58438 | Santa Cruz Biotechnology |
| PMEL17 | Mouse IgG1 | 1:100 | sc377325 | Santa Cruz Biotechnology |
| DCT | Mouse IgG2a | 1:100 | sc74439 | Santa Cruz Biotechnology |
| RhoA | Rabbit | 1:1,000 | 2117 | Cell Signaling Technology |
| RAC1/2/3 | Rabbit | 1:1,000 | 2465 | Cell Signaling Technology |
| CDC42 | Rabbit | 1:1,000 | 2462 | Cell Signaling Technology |
| F-actin | Mouse IgM | 1:100 | ab205 | Abcam |
| YAP | Rabbit | 1:1,000 | 8418 | Cell Signaling Technology |
| CCN1 | Rabbit | 1:1,000 | 14479 | Cell Signaling Technology |
| CCN2 | Rabbit | 1:1,000 | 86641 | Cell Signaling Technology |
| BRAF | Rabbit | 1:1,000 | 9433 | Cell Signaling Technology |
| Erk | Rabbit | 1:1,000 | 4695 | Cell Signaling Technology |
| p-Erk | Rabbit | 1:1,000 | 4370 | Cell Signaling Technology |
| p38 | Rabbit | 1:1,000 | 8690 | Cell Signaling Technology |
| p-p38 | Rabbit | 1:1,000 | 4511 | Cell Signaling Technology |
| JNK | Rabbit | 1:1,000 | 9252 | Cell Signaling Technology |
| p-JNK | Rabbit | 1:1,000 | 4668 | Cell Signaling Technology |
| c-Jun | Rabbit | 1:1,000 | 9165 | Cell Signaling Technology |
| p-c-Jun | Rabbit | 1:1,000 | 3270 | Cell Signaling Technology |
| c-Fos | Rabbit | 1:1,000 | 4384 | Cell Signaling Technology |
| p-c-Fos | Rabbit | 1:1,000 | 5348 | Cell Signaling Technology |
| JunB | Rabbit | 1:1,000 | 3753 | Cell Signaling Technology |
| p-JunB | Rabbit | 1:1,000 | 8053 | Cell Signaling Technology |
| JunD | Rabbit | 1:1,000 | 5000 | Cell Signaling Technology |
| GAPDH | Rabbit | 1:1,000 | 52118 | Cell Signaling Technology |

**Table S5.**
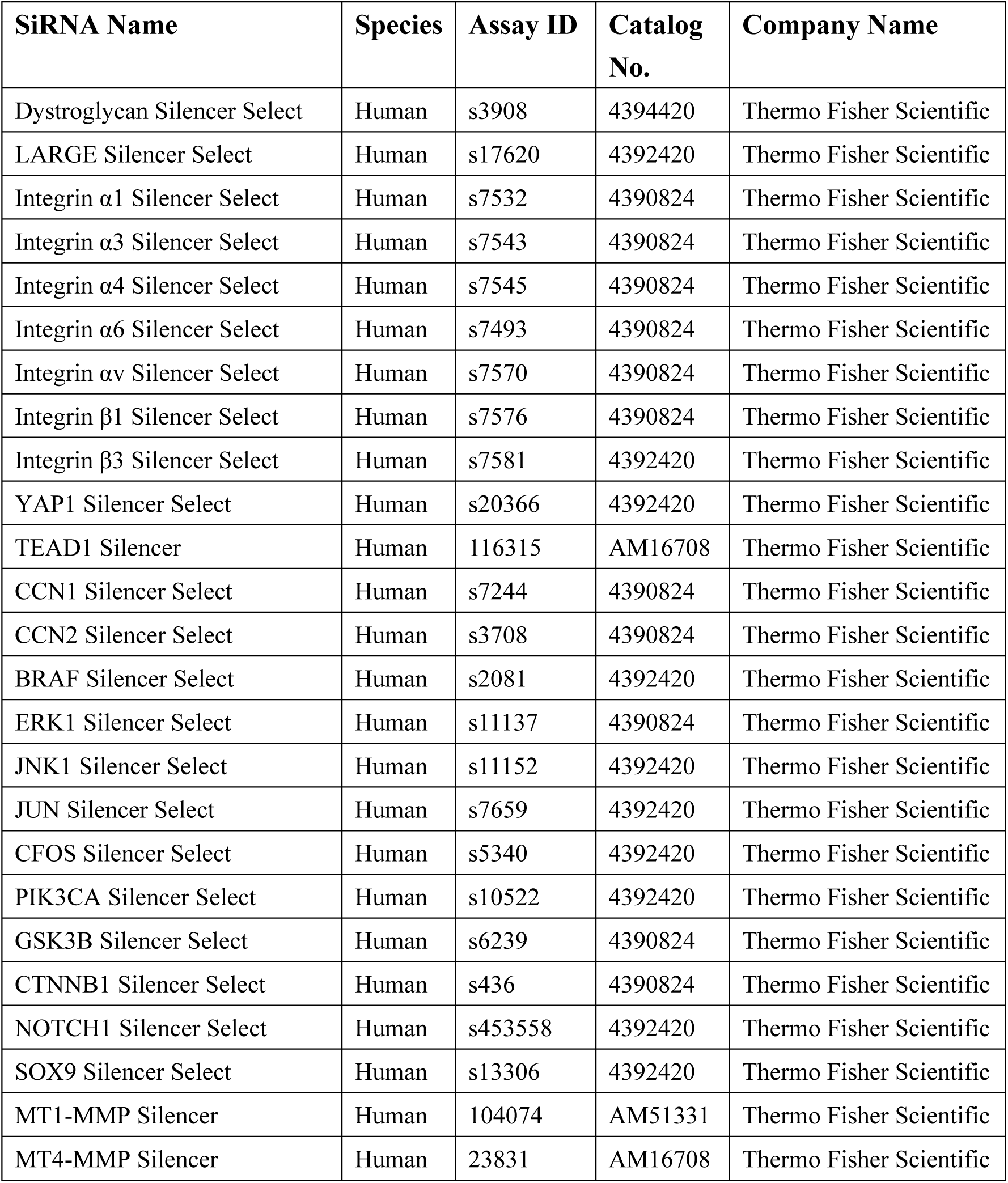
List of siRNAs used for siRNA-mediated knockdown in melanocytes.

**Table S6.** List of compounds used for *in vitro* drug treatments.

| **Compound Category** | **Compound Name** | **Treatment Concentration** | **Catalog No.** | **Company Name** |
| --- | --- | --- | --- | --- |
| Actin inhibitors | Cytochalasin D | 1 μM | 034-25881 | FUJIFILM Wako Pure Chemical Corporation |
|  | Latrunculin B | 250 nM | AG-CN2-0031-M001 | AdipoGen Life Sciences |
|  | Chaetoglobosin A | 500 nM | BVT-0092-M001 | AdipoGen Life Sciences |
| Actin activator | Jasplakinolide | 5 nM | AG-CN2-0037-C100 | AdipoGen Life Sciences |
| Rho/Rac/Cdc42 activator | Rho/Rac/Cdc42 Activator I | 2 μg/ml | CN04A | Cytoskeleton, Inc. |
| ROCK activator | Narciclasine | 40 nM | SML2805 | Sigma-Aldrich |
| YAP inhibitors | TED-347 | 150 nM | HY-125269 | MedchemExpress |
|  | Verteporfin | 300 nM | HY-B0146 | MedchemExpress |
| YAP activators | PY-60 | 10 μM | HY-141644 | MedchemExpress |
| | GA-017 | 10 μM | HY-147082 | MedchemExpress |
| JNK inhibitors | SP600125 | 50 μM | HY-12041 | MedchemExpress |
| | T-5224 | 25 μM | HY-12270 | MedchemExpress |
| JNK activator | Anisomycin | 100 nM | HY-18982 | MedchemExpress |

**Table S7.** List of compounds used for *in vivo* drug treatments.

| **Compound Category** | **Compound Name** | **Treatment Concentration** | **Catalog No.** | **Company Name** |
| --- | --- | --- | --- | --- |
| Actin inhibitor | Chaetoglobosin A | 500 μM | BVT-0092-M001 | AdipoGen Life Sciences |
| Actin activator | Jasplakinolide | 10 μM | AG-CN2-0037-C100 | AdipoGen Life Sciences |
| YAP inhibitor | Verteporfin | 500 μM | HY-B0146 | MedchemExpress |
| YAP activator | PY-60 | 500 μM | HY-141644 | MedchemExpress |
| JNK inhibitors | SP600125 | 500 μM | HY-12041 | MedchemExpress |
| JNK activator | Anisomycin | 200 μM | HY-18982 | MedchemExpress |

**Table S8.** List of compounds used for *ex vivo* drug treatments.

| **Compound Category** | **Compound Name** | **Treatment Concentration** | **Catalog No.** | **Company Name** |
| --- | --- | --- | --- | --- |
| Actin inhibitor | Latrunculin B | 2 μM | AG-CN2-0031-M001 | AdipoGen Life Sciences |
| | Chaetoglobosin A | 7.5 μM | BVT-0092-M001 | AdipoGen Life Sciences |
| Actin activator | Jasplakinolide | 50 nM | AG-CN2-0037-C100 | AdipoGen Life Sciences |
| YAP inhibitor | TED-347 | 1 μM | HY-125269 | MedchemExpress |
| | Verteporfin | 1 μM | HY-B0146 | MedchemExpress |
| YAP activator | PY-60 | 10 μM | HY-141644 | MedchemExpress |
| | GA-017 | 10 μM | HY-147082 | MedchemExpress |
| JNK inhibitors | SP600125 | 50 μM | HY-12041 | MedchemExpress |
| | T-5224 | 25 μM | HY-12270 | MedchemExpress |
| JNK activator | Anisomycin | 500 nM | HY-18982 | MedchemExpress |

